# Temporal Image Sandwiches Enable Link between Functional Data Analysis and Deep Learning for Single-Plant Cotton Senescence

**DOI:** 10.1101/2024.06.30.601428

**Authors:** Aaron J. DeSalvio, Alper Adak, Mustafa A. Arik, Nicholas R. Shepard, Serina M. DeSalvio, Seth C. Murray, Oriana García-Ramos, Himabindhu Badavath, David M. Stelly

## Abstract

Senescence is a highly ordered degenerative biological process that affects yield and quality in annuals and perennials. Images from 14 unoccupied aerial system (UAS, UAV, drone) flights captured the senescence window across two experiments while functional principal component analysis (FPCA) effectively reduced the dimensionality of temporal visual senescence ratings (VSRs) and two vegetation indices: RCC and TNDGR.

Convolutional neural networks (CNNs) trained on temporally concatenated, or “sandwiched,” UAS images of individual cotton plants (*Gossypium hirsutum* L.), allowed single-plant analysis (SPA). The first functional principal component scores (FPC1) served as the regression target across six CNN models (M1-M6).

Model performance was strongest for FPC1 scores from VSR (R^2^ = 0.857 and 0.886 for M1 and M4), strong for TNDGR (R^2^ = 0.743 and 0.745 for M3 and M6), and strong-to-moderate for RCC (R^2^ = 0.619 and 0.435 for M2 and M5), with deep learning attention of each model confirmed by activation of plant pixels within saliency maps.

Single-plant UAS image analysis across time enabled translatable implementations of high-throughput phenotyping by linking deep learning with functional data analysis (FDA). This has applications for fundamental plant biology, monitoring orchards or other spaced plantings, plant breeding, and genetic research.

## Introduction

### Senescence

Senescence encompasses the summation of gene-, cell-, tissue-, and organism-level changes leading to deterioration of biological function. Shuttling of assimilates from vegetative to reproductive organs in crop plants is a key feature of end-of-season senescence as it impacts harvest index of fruit, yield of grain or seed, nutrient composition, and efficiency of nutrient use (Gregersen et al., 2013). Structural changes to senescing cells occur in an ordered manner, with leaf senescence under nuclear control (Gan, 2003, Yoshida, 1962). Plant scientists frequently interpret senescence as a response to stress and factor it into selection decisions for breeding (Makanza et al., 2018), underscoring senescence as a quantitative selection metric that can simultaneously be analyzed as a time-series data modality.

Although it is grown commercially as an annual crop, *Gossypium* (cotton) maintains an indeterminate growth habit characteristic of perennials (Chen and Dong, 2016). As opposed to annuals, where nutrients are allocated to developing seeds, seasonal senescence in perennials is marked by nutrient shuttling to stems or roots, where they are reserved for growth in the next season (Woo et al., 2019). *G. tomentosum* is a wild allotetraploid species native to the Hawaiian Islands, where it is referred to as Ma’o (DeJoode and Wendel, 1992). Several *G. tomentosum* traits are desirable for introgression into cultivated cotton species, including its characteristic heat tolerance, resistance to pests and diseases such as fleahoppers, tarnished plant bug, bollworm, and boll rot (Saha et al., 2006), thrips and jassids (Hulse-Kemp et al., 2014), and for desirable agronomic traits including fiber quality, length, and fineness (Shim et al., 2018, Zhang et al., 2011). In cotton, late-season weather conditions influence senescence timing, with premature senescence and late boll maturity potentially conferring reductions in fiber quality and yield (Dong et al., 2006). Leaf senescence can be accelerated by extreme high or low temperatures, with high temperatures promoting an increase in chloroplast reactive oxygen species, thus damaging the chloroplast and photosynthesis-associated proteins, which ultimately impacts photosynthetic electron transfer (Ougham et al., 2008). Abiotic factors including drought, limited nutrients, oxidative stress via UV-B irradiation, and biotic factors such as plant pathogens and shading can each induce untimely senescence alone or in combination (Lim et al., 2003). The aging of many crop species demonstrates sensitivity to the source-sink ratio. Cotton senescence rate can be delayed by removal of fruiting branches (increasing the source-sink ratio by removing sink tissue) and accelerated by removing leaves (decreasing the ratio by removing source tissue) (Niu et al., 2007, Chen and Dong, 2016).

### High-throughput phenotyping analyses of senescence

Despite the importance of senescence as a trait, robust evaluations of large numbers of genotypes and/or genotype × environment combinations are complicated to evaluate across time (temporally) using repeated observations spanning the overall maturation period. Because the flowering habit of cotton is indeterminate, the maturation period can last weeks to months. This “phenotyping bottleneck” for senescence can be mitigated using unoccupied aerial systems (UAS, also known as UAVs or drones) for proximal sensing and modeling temporal phenotypes (Furbank and Tester, 2011) forming a basis of field-based high-throughput phenotyping (FHTP). An early application of greenhouse-based HTP proximal sensing showed that despite color distortion or blurring, empirically determined senescence scores could be obtained from images of Australian spring wheat (*Triticum aestivum*) and chickpea (*Cicer arietinum*) with moderate-to-strong positive correlations to visual scores (Cai et al., 2016). Lyu et al. (2017) reported a comprehensive greenhouse pipeline that dissects senescence at the single-leaf level using leaf tracking and principal component analysis (PCA) of empirically-determined senescence scores to cluster wild-type *Arabidopsis* and senescence mutants into groups with distinct temporal phenotypes. Visually scoring senescence in time-series UAS orthomosaics of maize hybrids, DeSalvio et al. (2022) identified quantitative phenotypic indicators of senescence progression through plot-based temporal vegetation indices. Each study demonstrated HTP (greenhouse- or field-based) can decrypt senescence quantitatively using spectral data. However, further methodological development and throughput of phenomics-based senescence characterization is required for plant biology, breeding, genetics, and commercial applications.

### Deep learning for plant phenotyping

Plant breeding programs depend on generational or yearly recording of phenotypic traits, many of which require time-consuming and labor-intensive collection methods. Accurately mapping connections between phenotype and genotype and ultimately saturating the phenome (Murray et al., 2022) requires methods that can shuttle broad classes of data through automated processing pipelines requiring little modification in each use case. Beginning with raw inputs such as images, deep learning methods function via a series of nonlinear layers that transform the raw inputs to slightly more abstract representations with each layer, ultimately amplifying “signal” from “noise” (LeCun et al., 2015). For example, early layers may detect basic features such as plant material vs. soil while more abstract layers can enable distinctions between patterns of plant and leaf pigmentation attributable to infections by different pathogens. Within the deep learning class of models, convolutional neural networks (CNNs) are suitable for image recognition and categorization, learning complex and nonlinear mappings from large example data sets (LeCun et al., 1998). CNNs are generally characterized by three types of neural layers: convolutional, pooling, and fully connected layers (Guo et al., 2016). During the forward stage of training, the input image, usually an RGB picture in the form of a 3-dimensional tensor (where X is height, Y is width, and Z is depth), is passed through each layer where the current weights and biases within each layer are applied, and the output (a prediction in the form of a class label or regression output) is subsequently compared with the ground measured “truth” value to calculate loss. After each convolutional layer, nonlinearity is often introduced using the Rectified Linear Unit (ReLU) function (Glorot et al., 2011) which also combats the vanishing gradient problem (Pawara et al., 2017). During the second stage of training, backpropagation entails iterative application of the chain rule to calculate the gradient of the loss function with respect to each parameter, with parameters updated based on these calculations (Guo et al., 2016). Fully connected layers generally employ dropout to avoid overfitting by randomly removing a percentage of nodes within the fully connected layer during training and backpropagation (Srivastava et al., 2014, Yoo, 2015, Pawara et al., 2017).

CNN applications in plant sciences include segmentation of overlapping field plants in maize (Guo et al., 2023), classification of soybean stress (Ghosal et al., 2018), and disease detection in bell pepper, potato, and tomato (Jung et al., 2023), wheat (Nigus et al., 2023), and across the PlantVillage data set, which includes 39 classes of plant leaves with varying diseases (Mohanty et al., 2016). Ubbens and Stavness (2017) demonstrated an early application of neural networks for leaf counting, classifying mutants, and plant age using primarily the International Plant Phenotyping Network Arabidopsis phenotyping data set (Minervini et al., 2014). In a regression context, Niu et al. (2024) reported the use of row- and grid-level temporally concatenated images where four RGB images, each with dimensions 32 × 32 × 3, were stacked along the third dimension (time) to produce 32 × 32 × 12 images (where 12 is calculated by 3 RGB channels per image × 4 images). With this input data structure, the CNN reported cotton yield values with lower mean absolute errors (MAE) and higher R^2^ values than AlexNet, ResNet, and CNN-LTSM models. This study highlighted that input data can be restructured such that time-series information is embedded in each input unit, thereby reducing the dimensionality of the data set down to the number of individual plots or in the case of single-plant analysis (SPA), individual plants.

No CNN method has yet been reported to enable temporal SPA of senescence in field-grown row crops. SPA would represent a paradigm shift from whole-plot analysis that is currently pervasive in crop experiments. Most SPA studies to date have focused on individual tree phenotyping and have been conducted manually without the benefit of CNNs. Novel SPA methods could enable increased statistical power without requiring increased land usage, refining the dissection of genotype × environment interactions at the single-plant level, and early-generation selection in plant breeding.

### Functional data analysis in plant sciences

Functional data analysis (FDA) is a statistical framework for the analysis and theory of data that vary over a continuum, such as repeated measures of a plant phenotypic trait (e.g., plant height, disease severity) across time. Functional principal component analysis (FPCA) enables dimensionality reduction and imputation when sparsity is present (Wang et al., 2016). FPCA can capture dominant modes of temporal variation embedded in noisy field data. To model temporal NGRDI vegetation index values of maize grown in irrigated and drought conditions, Adak et al. (2024) used FPCA to partition the spectral data based on the management conditions using the first and second functional principal components (FPC1 and FPC2) and to set the FPC1 and FPC2 values as response variables for quantitative trait locus (QTL) analysis. This pointed to well-known QTLs implicated in maize stress responses and growth regulation. In the current study, the concept of reducing dimensionality of temporal data via FPCA and using FPC1 scores as CNN regression target variables is explored in the context of temporal concatenation of single-plant senescence images, another dimensionality reduction technique. This complements methods proposed by Niu et al. (2024) for regression of cotton yield, however here, the regression target is a temporal phenotype (FPC1 senescence scores) instead of a yield value.

The methods proposed here serve to address the growing need for novel analysis techniques in dissecting the plant phenome and conducting targeted plant biology studies as well as breeding. The main objectives of this article were to: 1) perform dimensionality reduction of time-series single-plant senescence scores using visual and empirical (vegetation index-derived) senescence scoring methods; 2) train CNNs with temporally concatenated (“sandwiched”) single-plant images to learn mappings from input features to predict the target variable (FPC1 scores) of visual and empirical senescence scores; and 3) to perform a preliminary statistical analysis of time-series senescence data in cotton chromosome substitution lines (CSLs) and chromosome segment substitution lines (CSSLs).

## Materials and Methods

Fig. 1 depicts a graphical summary of the methods employed in this article. Descriptions of the methods used are listed in the order in which they appear in the figure.

**Figure 1.**
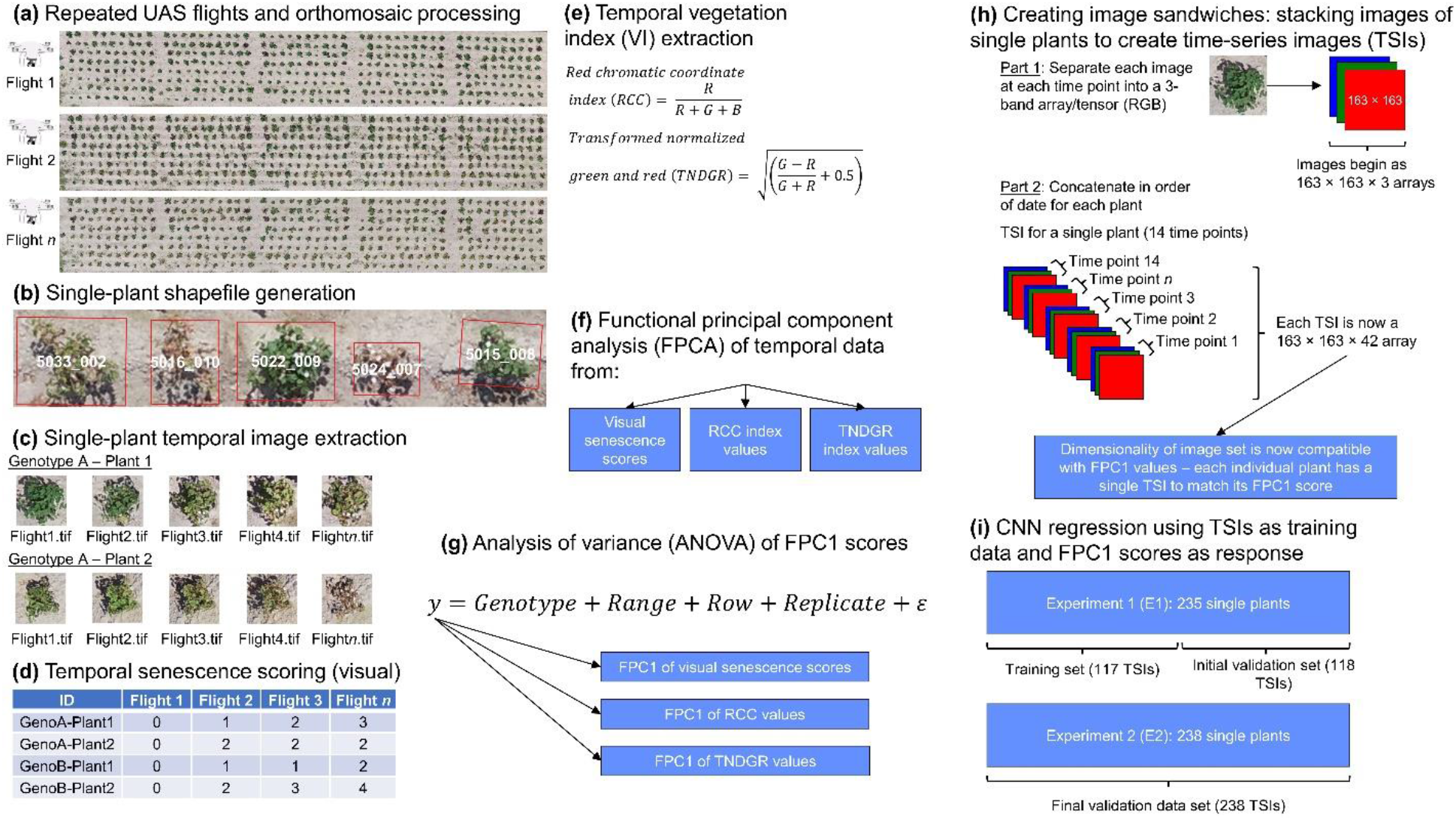
Graphical overview of data collection, visual and vegetation index-based senescence scoring, and preparation of data for CNN training and evaluation. (**a**) Orthomosaics from a representative sample of 3 of the 14 flights capturing the senescence window are shown in sequential order; (**b**) single-plant shapefiles were constructed and overlaid on each orthomosaic with minor adjustments to boundary boxes made as needed; (**c**) individual GeoTIFFs were extracted from each orthomosaic and converted to JPEGs; (**d**) plants were visually scored for senescence using images from (**c**); (**e**) empirical senescence scores were obtained by calculating vegetation indices for each image (**c**) using the FIELDimageR package; (**f**) functional principal component analysis (FPCA) was used to assess temporal data from visual senescence ratings (VSRs), RCC index values, and TNDGR index values; (**g**) the first functional principal component values, FPC1, were used in ANOVA; (**h**) time-series image (TSI), or “image sandwich” creation is outlined; (**i**) CNN data partitioning is presented, where optimal model hyperparameters were determined using 50% of TSIs from experiment 1 (E1) and evaluated on all images (unseen) from E2.

### Cotton germplasm and experimental design

A field experiment was conducted in College Station, TX, between April and September 2023 to evaluate upland cotton (*G. hirsutum*) BC5 backcross-inbred lines containing small portions of the genome of the wild Hawaiian cotton *G. tomentosum* (Nutt. ex Seem.), including 12 CSLs, 35 CSSLs, two upland lines (Texas Marker 1, TM-1), and a non-experimental "filler". Greenhouse-grown three-week old seedlings of 49 unique genotypes were mechanically space-transplanted into a randomized complete block design (RCBD) with 10 rows (*ca.* 200-cm spacing) of 10 plants (*ca.* 180-cm spacing), with outer rows and end hills serving as non-experimental "border", thus yielding 48 spaced transplants in each of the five blocks. Each genotype had approximately 5 replications with a total of 240 individual plants. Due to poor germination or other environmental causes, 5 plants perished early in the season, leaving 235 individual plants for time-series analysis. This experiment, hereafter abbreviated as E1, provided the training images for the CNN regression model. A concurrent field experiment (abbreviated E2) with 240 individual plants was conducted during the same timeframe and provided unseen validation images for CNN evaluation. Due to environmental causes or poor germination, 2 plants from E2 perished, leaving 238 available for analysis across 14 time points. Germplasm consisted of 27 genotypes (replicated approximately 9 times) of advanced backcross-inbred lines with a *G. hirsutum* background containing small introgressed portions of the *G. mustelinum* genome. As this experiment was primarily intended for seed increases and initial fiber quality characterization, it was not assessed via analysis of variance (ANOVA) and served strictly as validation data for CNN performance.

### High-throughput phenotyping of cotton experiments

After transplanting seedlings to the field, UAS flights were conducted two or three times each week totaling 46 flights across the growing season, of which 14 late-season flights were used to measure senescence occurring on: 24, 28, and 31 July; 4, 8, 11, 14, 16, 18, 21, 24, and 28 August; 1 and 5 September 2023. RGB images were stitched to produce orthomosaics (Methods **S1**) and single-plant shapefiles were created using the UAStools R package (Anderson and Murray, 2020) and adjusted where needed manually (Fig. **1a,b**; Methods **S2**).

### Single-plant temporal image extraction

Single-plant GeoTIFFs were extracted from orthomosaics and were converted to JPEGs (Fig. **1c**; Methods **S3**). In total, 235 and 238 images were obtained per flight, respectively, leading to a total data set size of 6,622 images (473 images per flight × 14 flights). Generally, the *RemSoil()* function within FIELDimageR (Matias et al., 2020) is advisable to limit the effects of soil on vegetation index calculations. However, inspection of images after soil removal indicated that senesced plant tissue was erroneously being removed and led to large holes in the images, likely due to similarity in color with soil, therefore this function was not employed for this study, and this limitation is acknowledged. To mitigate soil effects, shapefile bounding boxes were adjusted to crop closely around the borders of each single plant.

### Temporal senescence scoring of single-plant images (visual)

For each plant, 14 visual senescence ratings (VSRs) were assigned according to flight date and recorded in tabular format (6,622 total senescence scores). A scoring system with six levels was implemented, where 0 = 0% senescence (completely green), 1 = 20%, 2 = 40%, 3 = 60%, 4 = 80%, and 5 = 100% (completely dead) (Fig. **1d**). Examples of plants representing each score are shown in Fig. **2**. Three temporal phenotypes were observed: senescence progressed toward plant death (Fig. **2a**), stay-green occurred and vigor was maintained until the end of the season (Fig. **2b**), or plants presented an initial drop in vigor but displayed resilience and resurgence of vigor (Fig. **2c**). The transient display of intermediate senescence stages led to an imbalanced data set, which was dominated primarily by scores of 0, 1, 2, and 5, with notably less examples seen for 3 and 4, respectively (Table **1**). Due to this, and in keeping with previously published standard deep learning data augmentation practices (Ghosal et al., 2018), each image underwent a horizontal inversion and three rotations (clockwise) of 90°, 180°, and 270°. The total data set size consisted of 33,110 images (6,622 initial images + 6,622 × 4 augmentation methods) (Table **1**).

**Figure 2.**
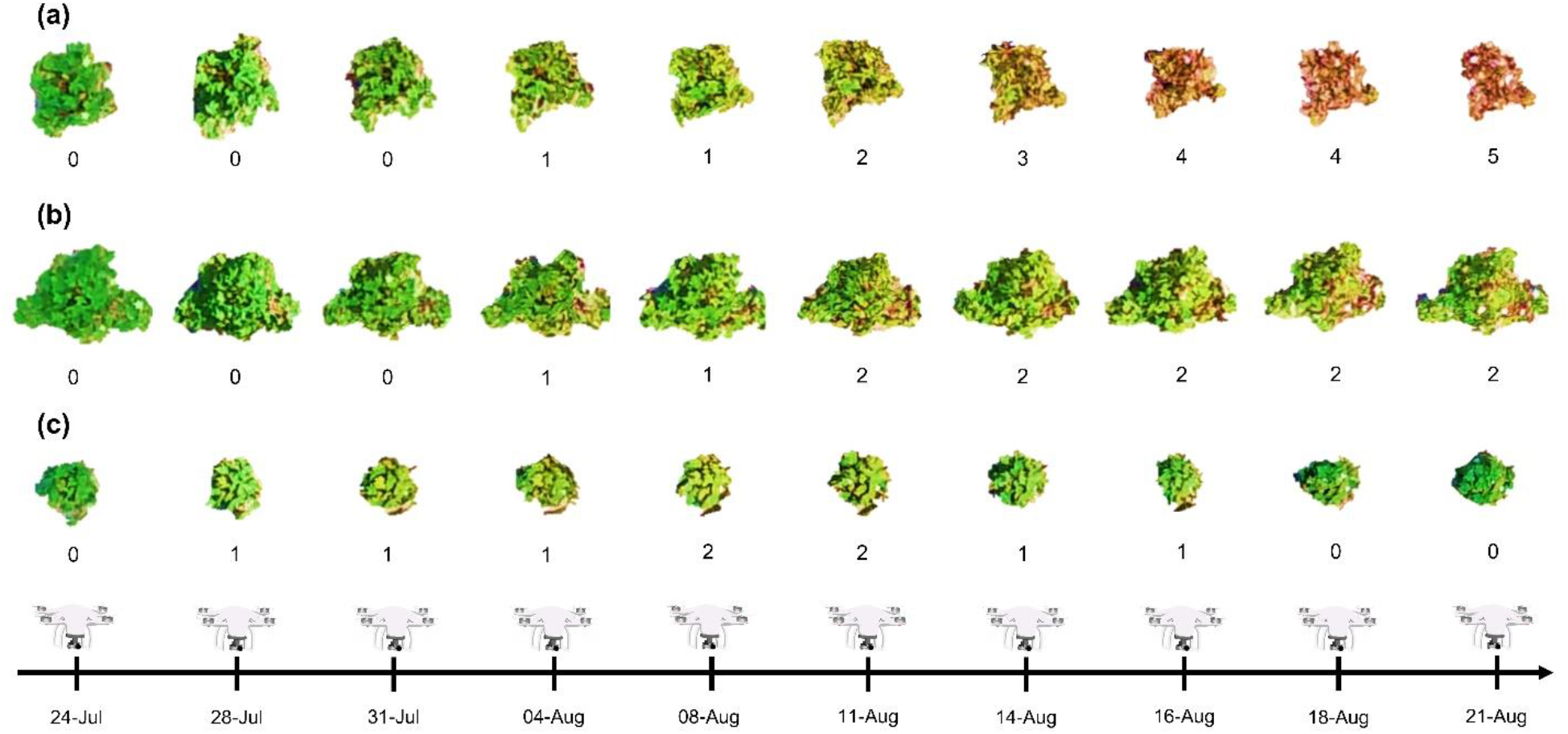
Three temporal phenotypes of senescence are shown for individual plants across 10 of the 14 flights selected for senescence scoring. In this figure only, contrast was enhanced, and soil was manually removed for illustration purposes, however this was not performed for CNN training and testing (raw images were used). (**a**) For this temporal phenotype, senescence progressed until permanent plant death. (**b**) Here, the stay-green phenomenon was observed, where the plant maintained intermediate greenness despite heat stress and drought. (**c**) Lastly, this temporal phenotype experienced an initial senescent episode but recovered and displayed late-season vigor.

**Table 1.** Distribution of senescence scores belonging to each category across experiment 1 (E1) and E2 revealed that many plants displayed either stay-green or a resurgence in vigor after an initial period of senescence. Data augmentation included images undergoing a horizontal flip and three clockwise rotations of 90°, 180°, and 270°.

| Visual Senescence Rating | 0 | 1 | 2 | 3 | 4 | 5 |
| --- | --- | --- | --- | --- | --- | --- |
| **E1** | 1,129 | 939 | 448 | 196 | 109 | 469 |
| **E2** | 1,303 | 1,096 | 474 | 127 | 59 | 273 |
| **Combined** | 2,432 | 2,035 | 922 | 323 | 168 | 742 |
| **E1 Augmented** | 5,645 | 4,695 | 2,240 | 980 | 545 | 2,345 |
| **E2 Augmented** | 6,515 | 5,480 | 2,370 | 635 | 295 | 1,365 |
| **Combined Augmented** | 12,160 | 10,175 | 4,610 | 1,615 | 840 | 3,710 |

### Temporal vegetation index (VI) extraction

Using the *fieldIndex()* function within FIELDimageR, 34 vegetation indices (VIs) were calculated for each plant at each time point (Fig. **1e**). Formulas for VIs are presented in **Supporting Information Table S1**. To identify potential candidates for empirical senescence scoring using VIs, each index was sequentially assessed for its Pearson correlation coefficient with visual scores using the *cor()* function in R. Two were selected (RCC and TNDGR) and the rest are not further described.

### Functional principal component analysis (FPCA) of temporal data and analysis of variance (ANOVA) of FPC1 scores

The fdapace R package (Wang et al., 2016, Chen et al., 2017) was used to perform both FPCA and prediction/imputation of values occurring during the senescence time grid for VSR, RCC index values, and TNDGR index values (Fig. **1f**). The *FPCA()* function within fdapace involves solving the integral FPCA expansion listed below (Karhunen, 1946, Loève, 1946):

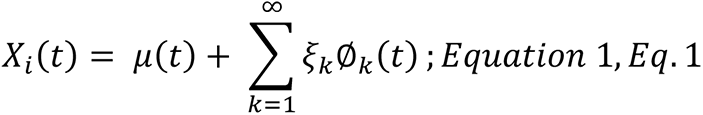

Here, 𝜇(𝑡) represents the mean function and indicates the average senescence progression at each time point 𝑡 (i.e., the date of each drone flight); 𝜉_𝑘_indicates the functional principal component scores for each function; ∅_𝑘_(𝑡) represent the eigenfunctions of 𝑘 functional principal components. To impute temporal values within the senescence time grid for VSRs, RCC, and TNDGR index values, the *predict()* function within fdapace was used.

FPCA was performed under two frameworks. The first framework sought to enable analysis of variance (ANOVA, Fig. **1g**) of temporal senescence. In this case, the first functional principal component scores (FPC1) from analyzing E1 data were used as the response variable. This framework used data from only E1, as ∼30% of plants within E2 had to be replaced with greenhouse backups several weeks after transplanting due to environmental causes. Therefore, E2 temporal scores were not evaluated using ANOVA. Since variance decomposition using ANOVA aims to partition genetic, environmental, and other effects from unexplained variation, only data from E1 was analyzed in this capacity.

The second framework was intended to prepare target variables for CNN regression. Pooled senescence data from E1 and E2 were assessed with FPCA. FPC1 scores from FPCA of combined E1 and E2 data were stored for CNN training and evaluation. The primary mode of temporal senescence progression (VSRs, RCC, or TNDGR) for each plant was captured by FPC1 and later served as the CNN target variable. The two FPCA frameworks produce unitless values for each eigenfunction (FPC1, FPC2, etc.), thus if E1 and E2 data were not pooled to obtain regression target values, it is possible that the ranges of FPC1 values could have differed between the two experiments (i.e., FPC1 values of 10 in E1 and E2 may not confer the same temporal phenotype).

To estimate genetic and field spatial variance components of FPC1 values for E1 VSRs, RCC, and TNDGR index values, ANOVA was performed via the lme4 R package (Bates et al., 2014) using Eq. **2**:

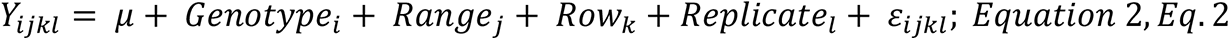

𝑌 is a vector of length 235 indicating each FPC1 value for observations of the 𝑖^𝑡ℎ^ genotype, in the 𝑗^𝑡ℎ^ range, 𝑘^𝑡ℎ^ row, and 𝑙^𝑡ℎ^ replication; 𝜇 indicates the experimental mean; 𝐺𝑒𝑛𝑜𝑡𝑦𝑝𝑒 indicates the effect of the 𝑖^𝑡ℎ^ genotype, with 𝑖 = 1, 2, … 49, where 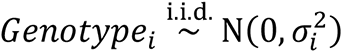; 𝑅𝑎𝑛𝑔𝑒 indicates the effect of the 𝑗^𝑡ℎ^ range, with 𝑗 = 1, 2, … , 40, where 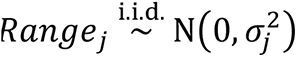; 𝑅𝑜𝑤 indicates the effect of the 𝑘^𝑡ℎ^ row, with 𝑘 = 1, 2, … , 6, where 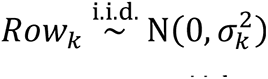; 𝑅𝑒𝑝𝑙𝑖𝑐𝑎𝑡𝑒 indicates the effect of the 𝑙^𝑡ℎ^ replication, with 𝑙 = 1, 2, … , 5, where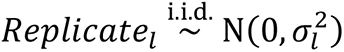; and 𝜀_𝑖𝑗𝑘𝑙_ indicates the residual error term that accounts for unexplained variability given by 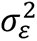 such that 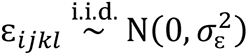.

Repeatability was calculated according to Eq. **3**:

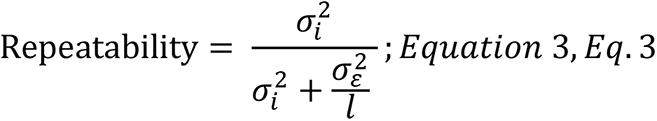

Here, 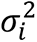 and 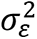 indicate the variance components of the effects of genotype and the error term, respectively, and 𝑙 is the number of replicates. In a minority of cases, replicates deviated from 5 due to seedling shortages or replacements of seedlings due to environmental conditions.

### Creating image sandwiches – stacking images of single plants to create time-series images (TSIs)

Images were imported into Python 3.11.5 using *iio.imread()* from imageio and resized to 163 × 163 × 3 with *tf.image.resize()* from TensorFlow, where 163 is the average image size (XY dimensions) and 3 indicates three RGB color channels. To create time-series images (TSIs) or “image sandwiches,” single-plant images were temporally concatenated using *np.concatenate()* (from NumPy) along the third dimension by flight date (Fig. **1h**). Each flight added three layers, resulting in TSIs of 163 × 163 × 42 pixels (3 channels × 14 flights). Each CNN experienced each plant as a temporally stacked image, with the FPC1 score as the regression target. The data set’s original dimensionality changed from 6,622 images of 163 × 163 × 3 (combined E1 and E2) to 473 TSIs of 163 × 163 × 42 pixels, corresponding to the number of single plants across E1 and E2. Post-augmentation, the total was 2,365 TSIs (473 initial + 1,892 augmented).

### CNN regression using TSIs as training data and FPC1 scores as response variables

Inspired by Zingaretti et al. (2020), hyperparameter optimization was undertaken using the Optuna Python package (Akiba et al., 2019) for each of the three regression scenarios: using TSIs to regress FPC1 scores of 1) VSRs, 2) RCC, and 3) TNDGR values. Optuna enables the user to create a “study,” whereby an objective function is defined with a parameter space through which Optuna systematically searches to achieve optimal performance at minimizing (in this case) or maximizing a target objective value across several “trials,” set at 250 in this study. Each trial was allowed to run for 50 epochs. Here, the study sought to find the hyperparameters that produced the lowest mean squared error (MSE). MSE was minimized because it is differentiable (its derivative can be calculated), which enables a smooth curve that gradient descent can travel to find the smallest error. Searchable hyperparameters are summarized in Table **2** and further details are provided in Methods **S4**. Six models underwent hyperparameter optimization, two each for VSRs, RCC, and TNDGR index values, with the only difference between them being the specification of the first dense layer always being set to ‘relu’ (M1-M3) or being searchable by Optuna (M4-6). Model names and associated traits are outlined in **Table 3**.

**Table 2.** The parameter space through which Optuna searched during hyperparameter optimization is presented.

| Hyperparameter | Type | Range/Choices | Notes |
| --- | --- | --- | --- |
| Dropout rate | Float | 0.0 to 0.5 | Uniformly distributed |
| Learning rate | Float | $1 \times 10^{-5}$ to $1 \times 10^{-2}$ | Logarithmically scaled |
| Regularization | Float | $1 \times 10^{-4}$ and $1 \times 10^{-2}$ | Logarithmically scaled |
| Activation function | Categorical | ‘relu’, ‘tanh’,<br>‘linear’ | Applies to Conv2D layers for first set of models; applies to Conv2D layers and first of two dense layers for second set of models |
| Dense neurons | Categorical | 128, 256, 512,<br>1024 | N/A |
| First kernel size | Categorical | 2, 3, 4, 5, 6, 7 | Size of kernel in first Conv2D layer |
| Initial filters | Categorical | 16, 32 | Number of filters in first Conv2D layer |
| Number of Conv2D layers | Integer | 1 to 5 | N/A |

**Table 3.**
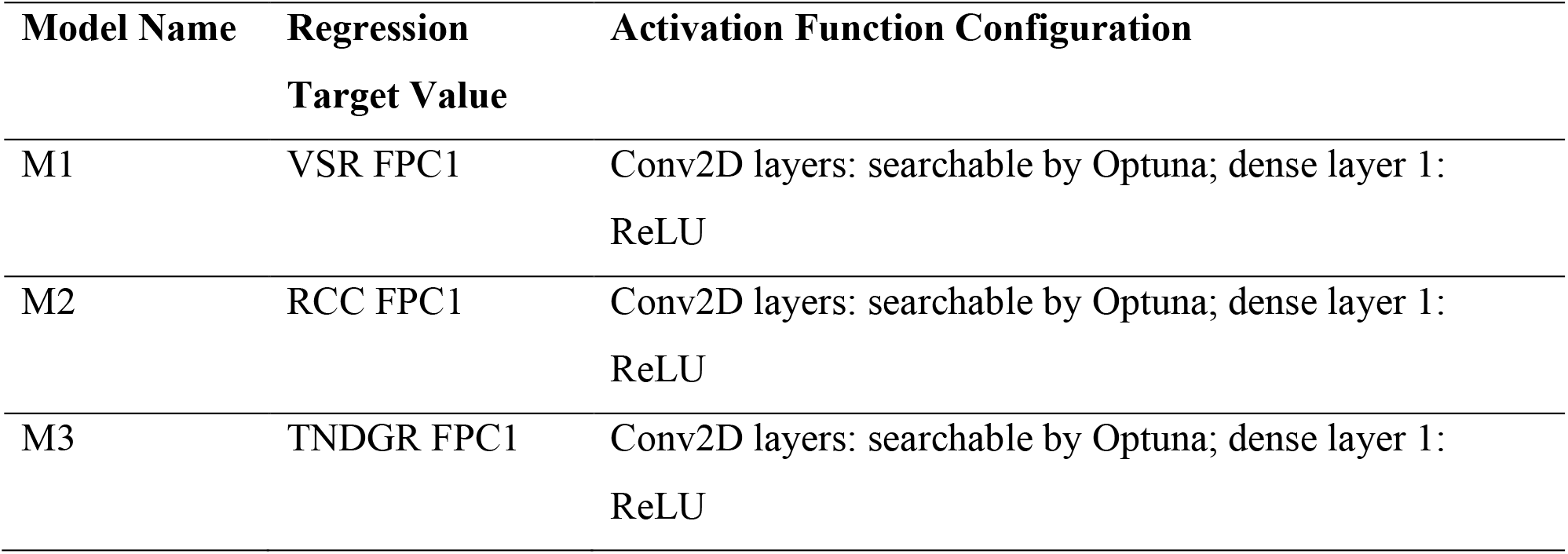
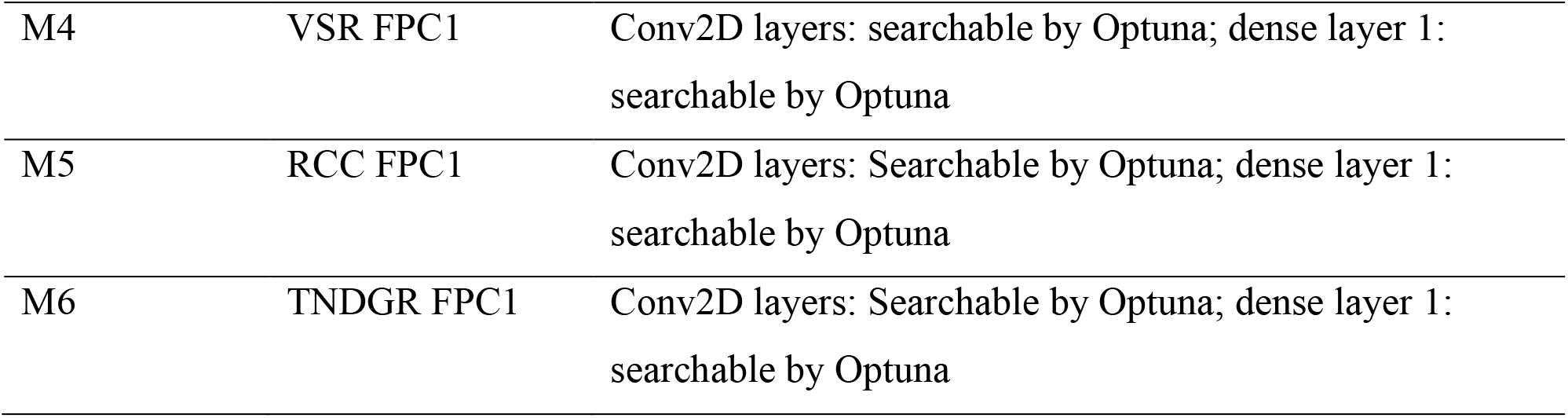
CNN model names, regression targets, and notes explaining the difference between the first set of models (M1-M3) and the second set (M4-M6). The regression targets are FPC1 scores of temporal senescence derived from pooled data from experiments E1 and E2 for: visual senescence ratings (VSRs), RCC, or TNDGR vegetation index values.

| Model Name | Regression Target Value | Activation Function Configuration |
| --- | --- | --- |
| M1 | VSR FPC1 | Conv2D layers: searchable by Optuna; dense layer 1: ReLU |
| M2 | RCC FPC1 | Conv2D layers: searchable by Optuna; dense layer 1: ReLU |
| M3 | TNDGR FPC1 | Conv2D layers: searchable by Optuna; dense layer 1: ReLU |
| M4 | VSR FPC1 | Conv2D layers: searchable by Optuna; dense layer 1:<br>searchable by Optuna |
| M5 | RCC FPC1 | Conv2D layers: Searchable by Optuna; dense layer 1:<br>searchable by Optuna |
| M6 | TNDGR FPC1 | Conv2D layers: Searchable by Optuna; dense layer 1:<br>searchable by Optuna |

### Model training and evaluation

Model development used TSIs from E1, whereas E2 served as an unseen validation set to avoid overfitting and assess general usability for different cotton germplasm. The same 50% of TSIs from E1 were used for training and Optuna hyperparameter optimization across six CNN models (Fig. **1i**), while the remaining 50% of E1 TSIs served as an initial validation set. Post-augmentation, the training set totaled 585 TSIs (117 initial + 468 augmented) and the validation set 590 TSIs (118 initial + 472 augmented), bringing E1’s total to 1,175 TSIs. Setting the randomization seed ensured consistent TSIs for all six Optuna studies, enabling direct model comparisons. Optimal hyperparameters were then used for CNN regression on the full E2 validation set. Final evaluation used 80% of the 1,175 TSIs from E1 for training and all 1,190 TSIs from E2 (238 initial + 952 augmented) for validation. In each replication, 940 TSIs from E1 served as training (0.8 × 1,175 = 940), with a randomization seed specified for repeatable splits. Metrics saved for each epoch included loss, validation loss, MAE, and validation MAE. R and R^2^ were calculated using R’s *cor()* function (Pearson correlation) to assess correlation between actual and predicted FPC1 values. MSE, RMSE, and MAPE were calculated using the Metrics R package. Hyperparameter importances were determined with functional ANOVA (fANOVA) using the *plot_param_importances()* function.

Models were also assessed through ANOVA of FPC1 scores output by the top performing CNNs for each senescence scoring metric to determine variance components (Eq. **2**) and repeatability of predicted FPC1 values (Eq. **3**). Using the optimal hyperparameters determined by Optuna, CNN regression of E1 FPC1 values was performed using the same train/test split that was used for the hyperparameter optimization trials, where a 50/50 train/test split was enacted with an internal 20% validation set. After performing 25 replications of CNN regression using a random 50/50 train/test split within each replication, predicted FPC1 values for each E1 plant were averaged, producing a vector of 235 FPC1 values that were subsequently assessed through ANOVA.

### Model explainability

In keeping with the precedent set by Ghosal et al. (2018) for plant stress phenotyping and initially by Erhan et al. (2009) and Simonyan et al. (2013), saliency maps were generated for each model using a representative TSI from a stay-green and rapid senescence plant (**Supporting Information Fig. S1-6**). Map generation is outlined in Methods **S5**.

## Results

### Correlations between vegetation indices and visual senescence ratings

In total, 34 vegetation indices (VIs) were assessed for their correlations with VSRs (Fig. **3a**) using all available data from E1 and E2, with formulas for each VI presented in **Table S1**. The highest positively-correlated index with manual senescence scores was the red chromatic coordinate index (RCC) (Woebbecke et al., 1995) with *R* = 0.75 (Fig. **3b**). However, the score with the highest magnitude of correlation, though negative, was the transformed normalized green and red index (TNDGR) (Tucker, 1979) with *R* = -0.81 (Fig. **3c**).

**Figure 3.**
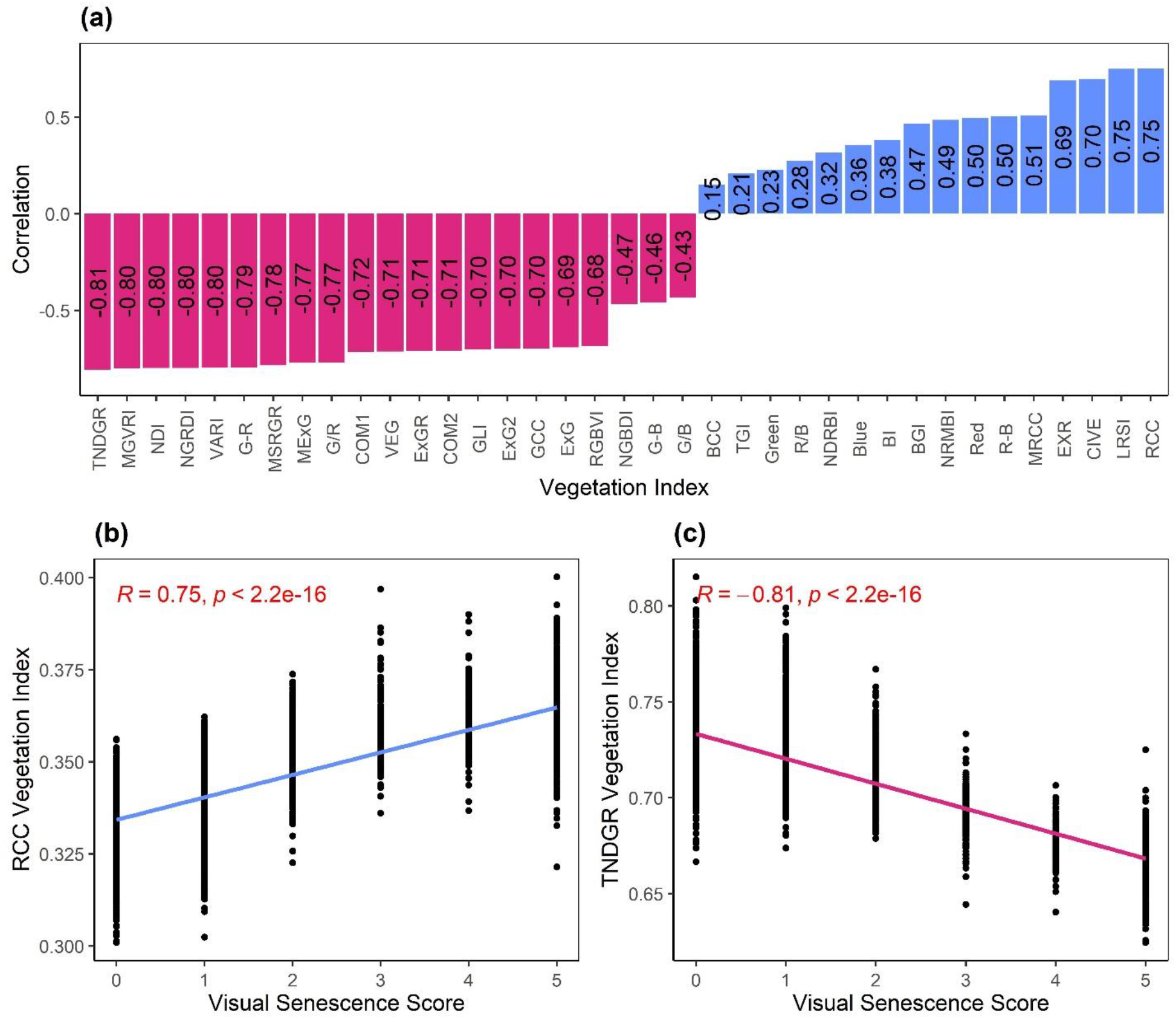
(**a**) Correlations of all 34 vegetation indices (VIs) with visual senescence ratings (VSRs). (**b**) The red chromatic coordinate index (RCC) displayed the highest positive correlation with VSRs, while TNDGR (**c**) displayed the highest magnitude of correlation with VSRs.

### Functional principal component analysis

FPCA was performed to synthesize a single value from 14 temporal measures for each plant under two frameworks (*Materials and Methods*). When FPCA was performed using data from E1 only, FPC1 and FPC2 explained 89.1% and 7.3% for visual scores, 83.5% and 10.1% for RCC, and 86.1% and 8.9% for TNDGR, respectively. For VSRs using pooled data from E1 and E2, FPC1 explained 90.2% of the temporal variation (Fig. **4a**), with FPC1 explaining 83.5% (Fig. **4b**) and 86.9% (Fig. **4c**) of the temporal variation for RCC and TNDGR, respectively. In the VSR functional principal component plot, manual inspection of the 14 images associated with each point (which denotes a single plant) indicated that generally, plants with an FPC1 score exceeding 10 could be classified as having a “rapid senescence” phenotype (Fig. **2a**; Fig. **4a**), while those with FPC1 scores below 10 corresponded to “stay-green” or resilient phenotypes noted in Fig. **2b** and **2c**. A distinct cluster of plants exhibiting rapid senescence is visible on the right side of the VSR FPCA plot (Fig. **4a**). Clustering was less pronounced for RCC (Fig. **4b**), but TNDGR displayed moderate partitioning of temporal phenotypes on the left side of the plot (Fig. **4c**).

**Figure 4.**
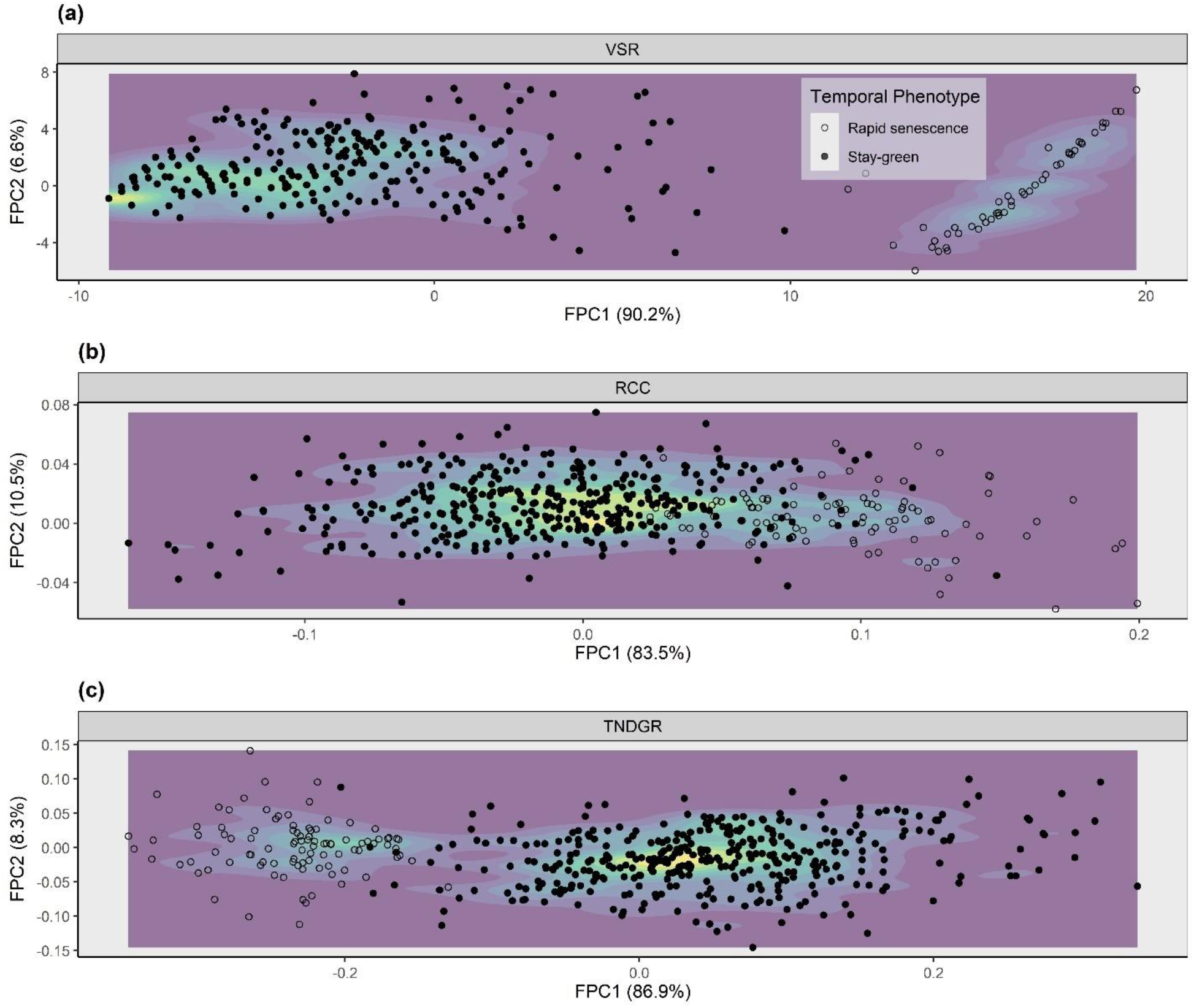
Plots of the first two functional principal components (FPC1 and FPC2) are presented from FPCA of combined senescence-tracking metrics from experiment 1 (E1) and E2 based on (**a**) visual senescence ratings (VSR), (**b**) RCC, and (**c**) TNDGR vegetation index values. Open circles indicate plants classified as rapidly senescing based on VSR FPC1 > 10.

### Temporal senescence trajectories of single plants as predicted by FPCA

Temporal trajectories of predicted FPCA scores for VSRs of plants within E1 (Fig. **5a**) revealed a similar pattern of separation as the FPCA plots (Fig. **4a**). Rapidly senescing single plants displayed a prominent rise in senescence between 110 and 130 days after transplanting (DAT). Six genotypes from E1 were chosen to illustrate potential genotypic differences in temporal senescence (Fig. **5b**), in which four of five replicates within genotypes 5002, 5012, and 5021 underwent rapid senescence while all replicates from genotypes 5011, 5036, and 5038 demonstrated varying degrees of stay-green.

**Figure 5.**
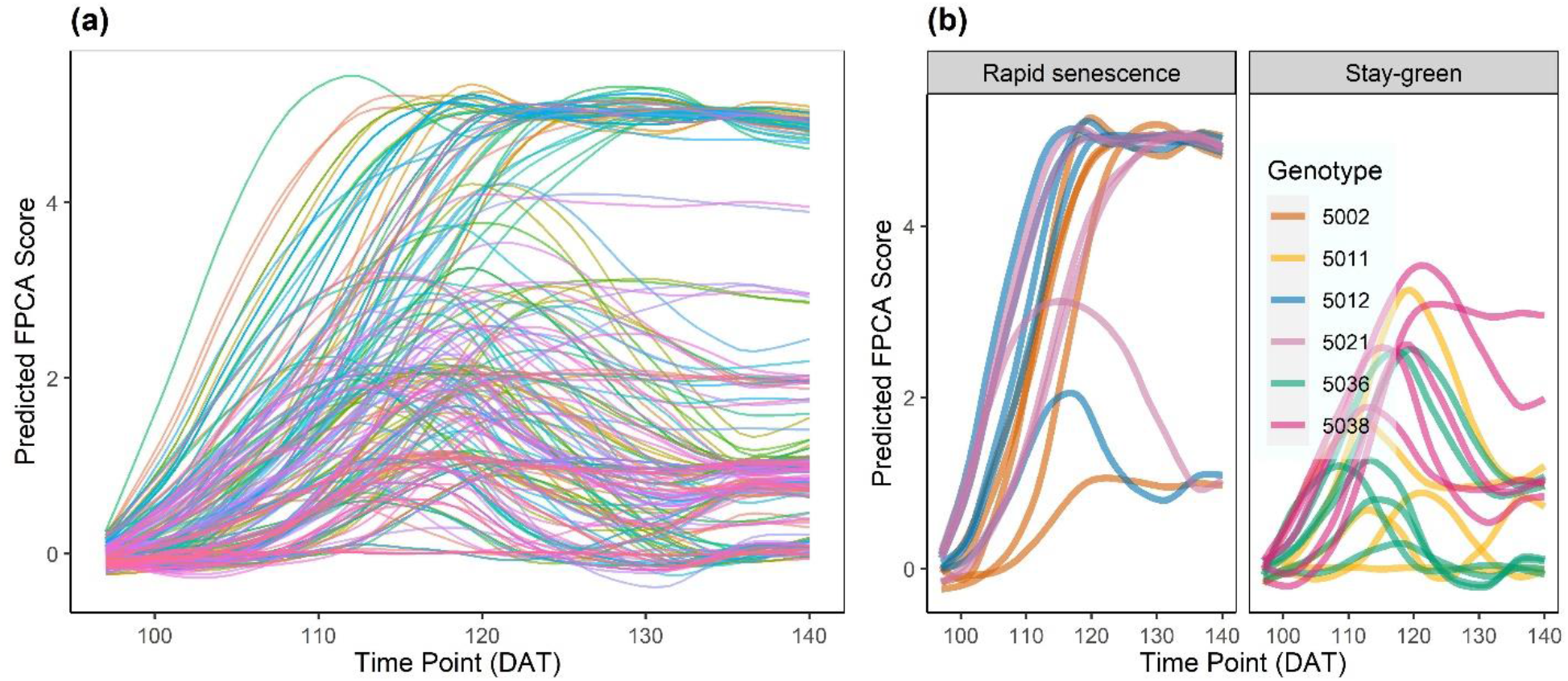
Predicted FPCA scores of visual senescence ratings are presented for (**a**) all 235 single plants in experiment 1 (E1) and for (**b**) six selected genotypes and their respective replicates.

### Analysis of variance (ANOVA) of temporal senescence phenotypes with FPC1 scores

ANOVA is an objective way to compare the consistency and precision of different measures between replicates. Results of ANOVA with Eq. **2** (E1 data only) indicated the highest percentage of variation captured by genotype (36.8%) was produced by setting FPC1 scores of TNDGR as the response (Fig. **6c**), followed by FPC1 scores of VSRs (31.4%, Fig. **6a**) and RCC (29.9%, Fig. **6b**). RCC reported the highest degree of field spatial variation, with 21.1% of variability in FPC1 scores attributable to row effects and 9.6% attributable to replicate effects. Visual scores demonstrated the highest degree of unexplained error variation at 55.3%. Repeatability, as calculated by Eq. **3**, was highest for TNDGR (0.80), while R^2^ was highest in RCC (0.61).

**Figure 6.**
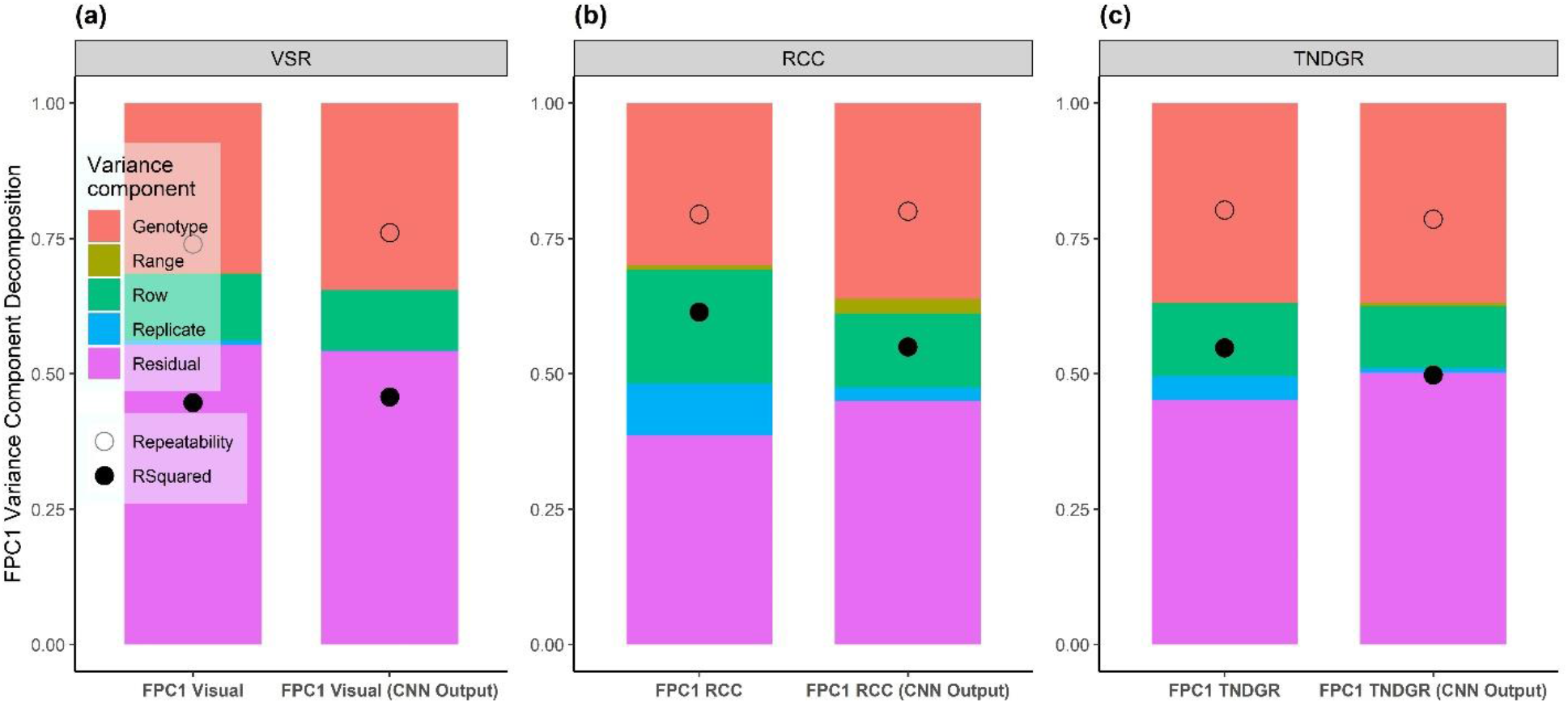
Graphical depiction of analysis of variance (ANOVA) for setting FPC1 scores of (**a**) VSRs, (**b**) RCC, and (**c**) TNDGR vegetation indices as response variables for experiment 1 (E1). Open circles indicate repeatability values as calculated by **Eq. 3** and closed circles denote R^2^ values for each trait. Within each pane, the left side reports ANOVA of FPC1 scores calculated directly from the data, whereas the right side reports ANOVA of CNN regression output of the top- performing model for each senescence metric. CNN training and testing here occurred within E1.

Variance decomposition of FPC1 values from the top-performing CNN regression models (regressing E1 data only) within each senescence measure indicated that genotypic variance was captured in a greater proportion for visual scores and RCC, improving over ANOVA results from actual FPC1 scores to 34.5% (Fig. **6a**), 36.2% (Fig. **6b**), respectively, while staying at 36.8% (Fig. **6c**) for TNDGR. Repeatability using CNN models improved 2.9% for visual scores, remained constant for RCC, and decreased 2.1% for TNDGR while R^2^ increased 2.4% for visual scores and decreased 10.5% and 9.1% for RCC and TNDGR, respectively.

### Hyperparameter optimization

The hyperparameters empirically determined by Optuna to perform best for regression of each temporal senescence data set are presented in **Tables S2-4**, where CNN model structure summaries detailing the output shapes and parameter numbers at each layer are also detailed (**Tables S5-10**). ReLU was chosen as the activation function for all models except M4, where hyperbolic tangent was selected. Except for M3 and M6 in TNDGR, the learning rate exerted the highest variable importance based on functional ANOVA results (Fig. **7a**), explaining a maximum of 94% of variation in M1 (visual score). Across the 250 trials, results of each trial’s performance at attempting to minimize the MSE are reported, with local minima indicated by a pink/red line that progresses stepwise as a new minimum is found (Fig. **7b**).

**Figure 7.**
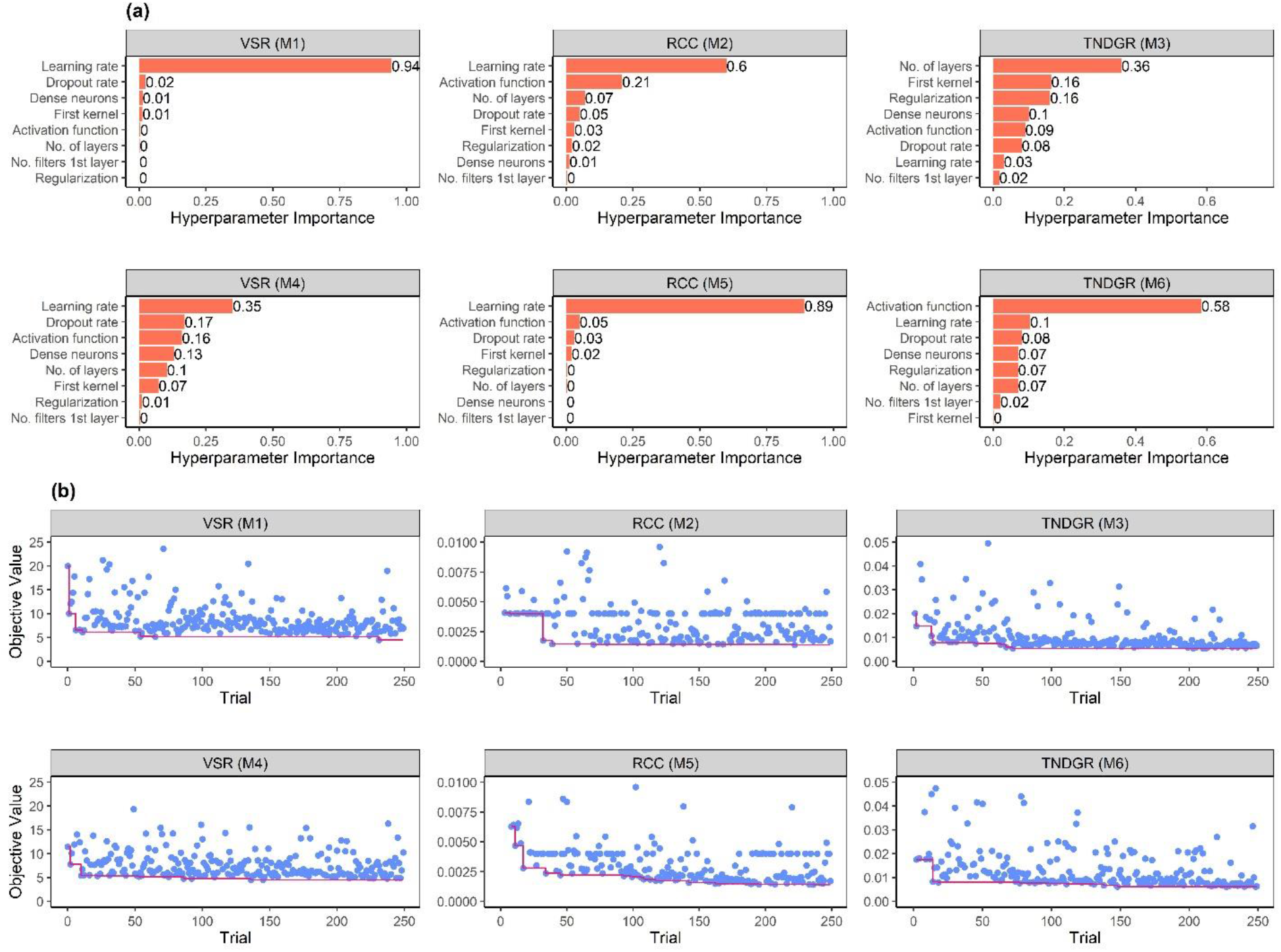
Assessment of models M1-6 according to hyperparameter variable importance scores determined by functional analysis of variance (fANOVA) using the Optuna Python package. M1, M2, and M3 variable importances are presented in (**a**), with M4, M5, and M6 importance scores presented in (**b**). Each Optuna trial’s mean squared error (MSE) value is indicated by a blue circle in (**c**) and (**d**). M1, M2, and M3 trial results are given in (**c**) while (**d**) indicates results from M4, M5, and M6. The pink/red lines at the bottom of (**c**) and (**d**) indicate local minima (improvements) as they were discovered across the 250 trials. Y-axes are identical within each trait for ease of comparison between models (visual score - M1 and M4, RCC - M2 and M5, and TNDGR - M3 and M6), but within each trait, some outlier MSE values are not visible as the Y-axes were truncated for visual clarity of the majority trial results.

### CNN regression model training metrics

Mean values from 25 replications of evaluating M1-6 with unseen TSIs (the entirety of the E2 TSI set) are reported across 50 epochs of training within each replication with loss (Fig. **8a**) and MAE (Fig. **8b**). Loss and MAE scales are different for each of the three temporal senescence metrics as noted by the differences in Y-axis value ranges.

**Figure 8.**
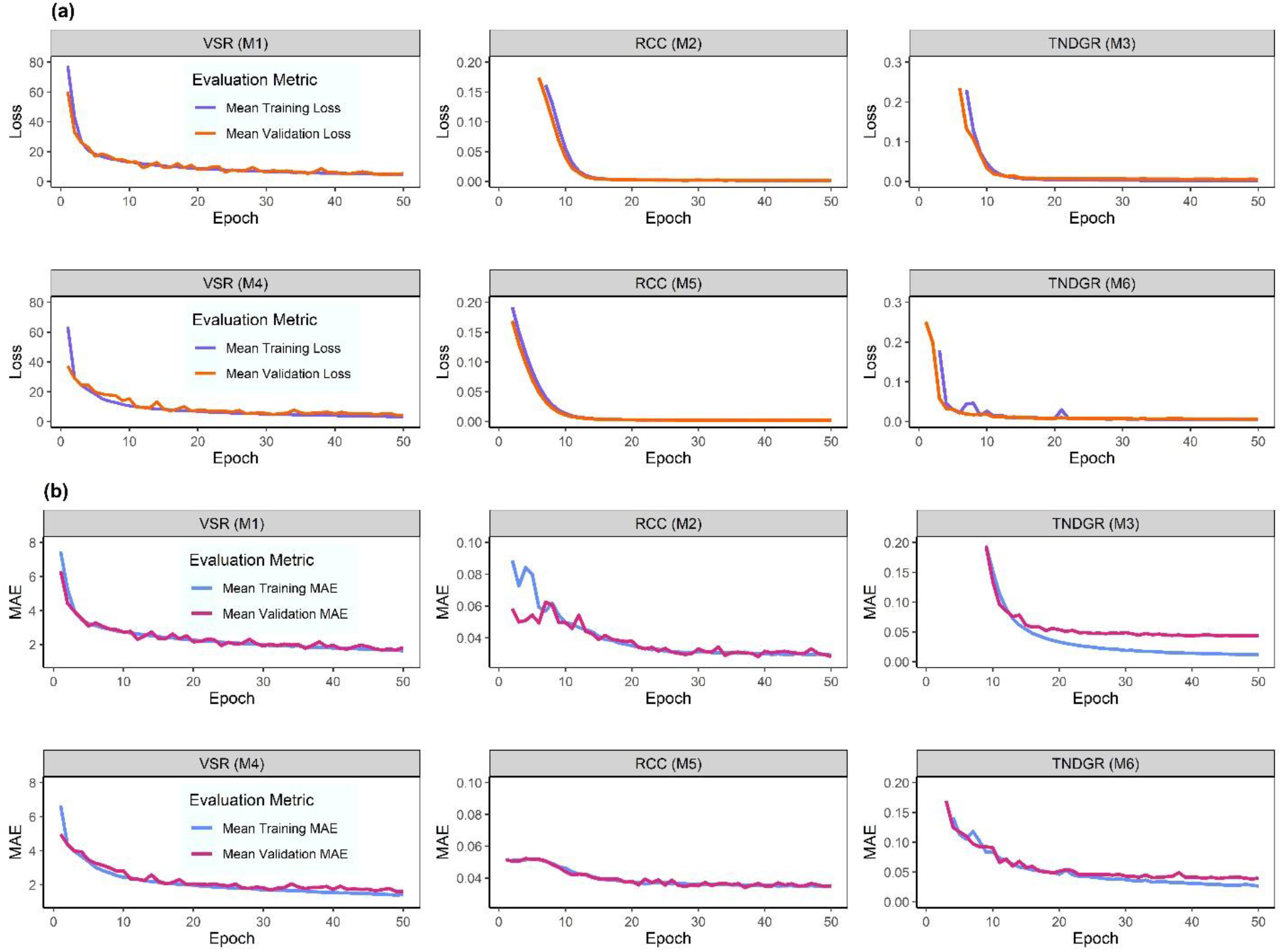
All six CNNs (M1-6) were evaluated with 25 replications by using Optuna-derived hyperparameters. In each replication, models were trained using 80% of time-series images (TSIs) from E1 (with an internal 20% validation image set) and were tasked with outputting FPC1 values from all E2 images, which represented unseen validation TSIs. Loss values (averaged from 25 replications) are presented in (**a**). Mean absolute error (MAE) values, again averaged across 25 replications, are given in (**b**). For RCC and TNDGR results, some initial loss and MAE values were outliers and hence the Y-axes have been truncated for visual clarity.

### CNN regression model performance

Performance metrics for all six CNN regression models (M1-6) across 25 replications are presented in Table **4**. All metrics refer to regression results from unseen validation images (the entire E2 data set). R and R^2^ were assessed using the *cor()* function in R, while root mean squared error (RMSE), mean absolute error (MAE), mean squared error (MSE), and mean absolute percentage error (MAPE) were calculated using actual and predicted values within respective functions from the Metrics R package. FPC1 values of VSRs were predicted by M1 with an R^2^ value of 0.857 and a mean absolute percentage error (MAPE) of 1.12%, while M4 performed the strongest of all models with an R^2^ of 0.886. Model performance for RCC was strong for M2 (R^2^ = 0.619) and moderate for M5 (R^2^ = 0.435), both with large MAPE values. Performance for TNDGR was strong with both M3 and M6 exceeding R^2^ = 0.74.

**Table 4.**
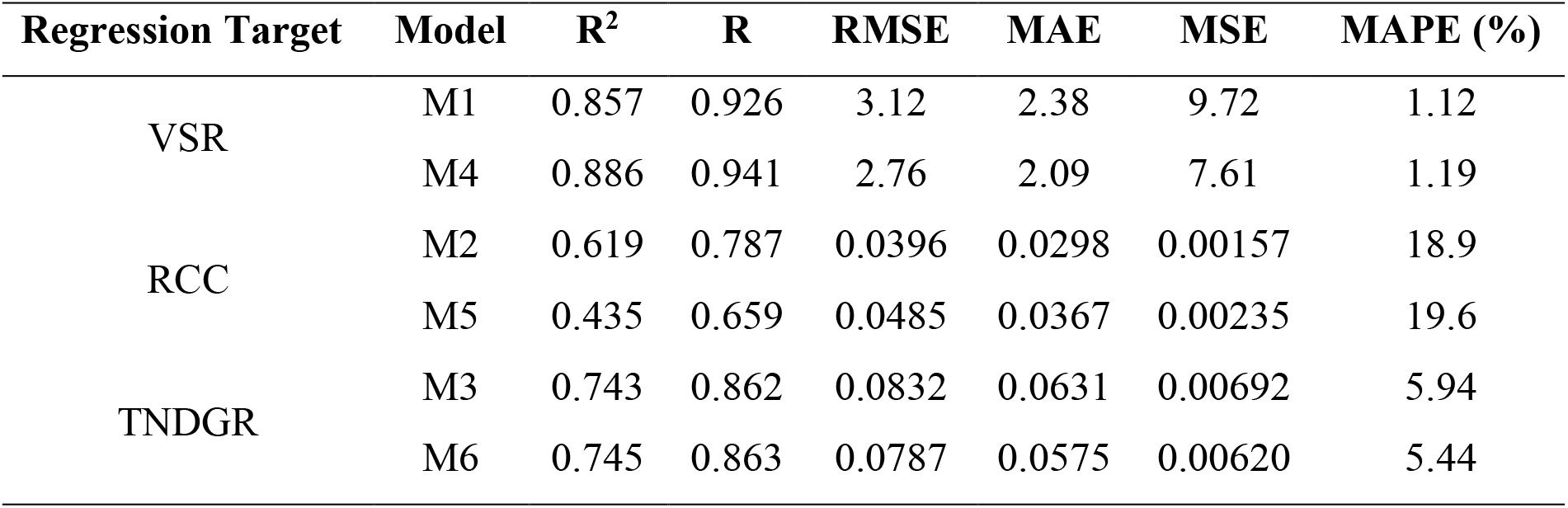
Model performance metrics for M1-6 calculated using actual and predicted values from CNN regression. Data are grouped according to regression target variables due to the differences in scale associated with each target that affect the interpretation of root mean squared error (RMSE), mean absolute error (MAE), mean squared error (MSE), and MAPE.

| Regression Target | Model | $R^2$ | R | RMSE | MAE | MSE | MAPE (%) |
| --- | --- | --- | --- | --- | --- | --- | --- |
| VSR | M1 | 0.857 | 0.926 | 3.12 | 2.38 | 9.72 | 1.12 |
|  | M4 | 0.886 | 0.941 | 2.76 | 2.09 | 7.61 | 1.19 |
| RCC | M2 | 0.619 | 0.787 | 0.0396 | 0.0298 | 0.00157 | 18.9 |
|  | M5 | 0.435 | 0.659 | 0.0485 | 0.0367 | 0.00235 | 19.6 |
| TNDGR | M3 | 0.743 | 0.862 | 0.0832 | 0.0631 | 0.00692 | 5.94 |
|  | M6 | 0.745 | 0.863 | 0.0787 | 0.0575 | 0.00620 | 5.44 |

### Model explainability

Examination of activation maps from the stay-green (Fig. **9a**) and rapid senescence (Fig. **9b**) temporal phenotypes from M4 revealed that the highest activation region (red) was centered on the cotton plant in both cases. This indicates the model was relying on plant pixels when outputting regression target values across the entire TSI. Regions of lesser yet still noteworthy activation were observed at the edges of many of the TSIs for saliency maps generated from M2/5 and M3/6, potentially indicating heightened sensitivity of VIs to late-season weed pressure that occurred in both E1 and E2 (Fig. **S3-6**).

**Figure 9.**
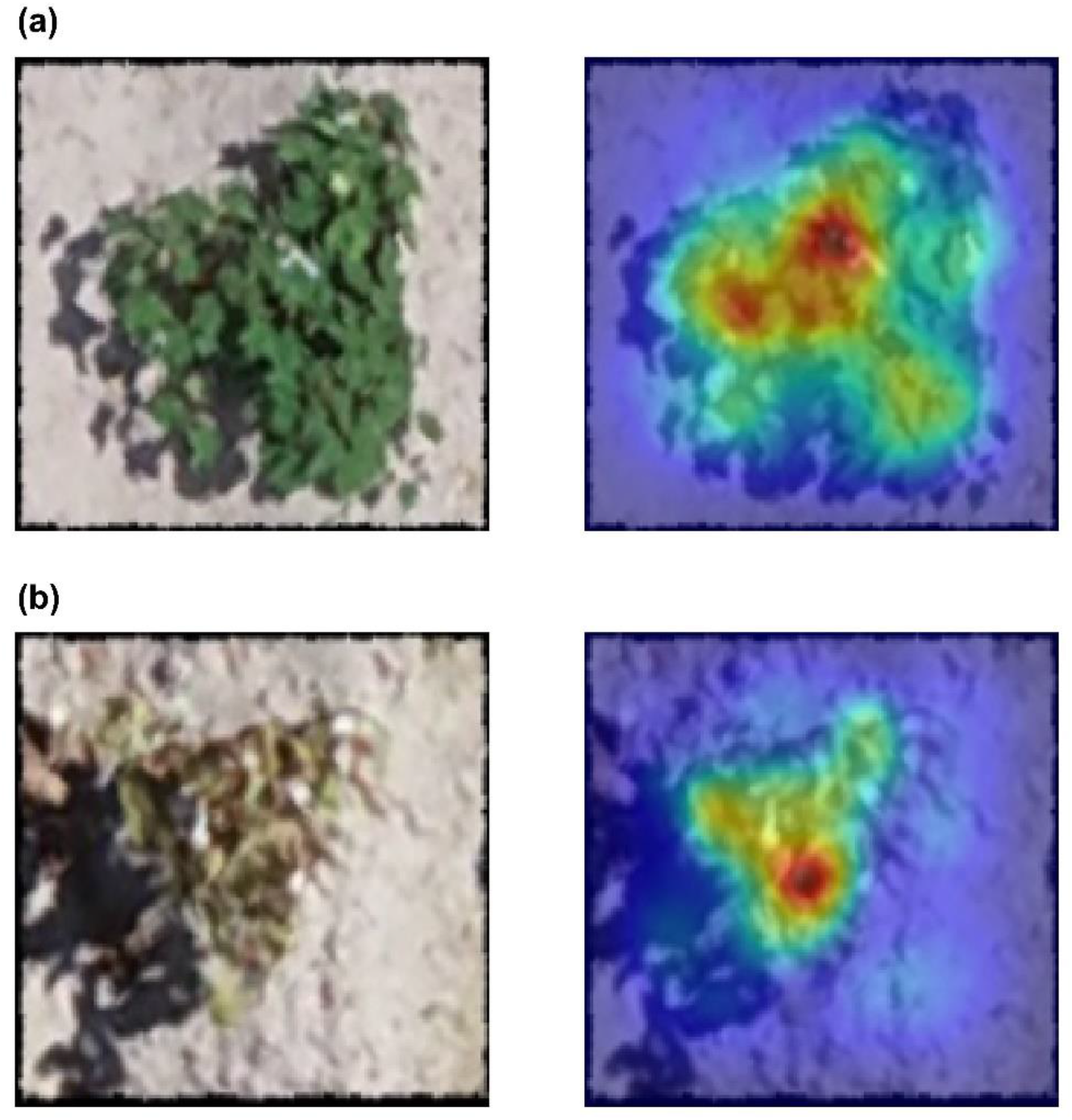
Saliency/activation maps from model M4 for two plants, one with (**a**) a stay-green phenotype and the other with (**b**) a rapid senescence phenotype. Both plants shown were from the E2 validation set that were not used in model training. Colors closer to red indicate regions of strong activation.

## Discussion

Leaf senescence in higher plants such as cotton (*Gossypium hirsutum* L.) entails communication between pathways involved in environmental sensing, hormones, and circadian rhythms (Gan and Amasino, 1997, Lim et al., 2007, Lee et al., 2021). Senescence is inherently a functional trait since it progresses along a continuum, thus, the amount of data collected for individuals in a senescence study depends on the time of the researcher and the instrumentation throughput. Elucidating mechanisms underlying senescence has agronomic and biological implications for a myriad of crops, particularly due to its implications for nutrient shuttling from source to sink tissue, with nitrogen being the predominantly remobilized element (Gregersen, 2011, Gregersen et al., 2013, Havé et al., 2017). Identification of crops of any type whose senescence trajectories are fine-tuned to a unique growing environment requires selection principles rooted in the temporal trajectory of senescence. This study demonstrated that field-based high-throughput phenotyping (FHTP) of spaced single plants using unoccupied aerial systems (UAS) can provide time-series spectral data that can be paired with functional data analysis (FDA) techniques such as functional principal component analysis (FPCA) to improve biological signals (genetic variance and repeatability) while reducing the dimensionality of time-series data (Fig. **4**). This approach simplifies decisions for future experimentation or can enable fine-tuning of selections for environmentally adapted material. Decoding senescence has yielded success in basic biology, revealing the tight regulation of the senescent transcriptome (Woo et al., 2016), the interplay between plant hormones such as abscisic acid, senescence, and drought resistance (Zhao et al., 2016), and the activity of time-evolving gene regulation at presenescent stages (Kim et al., 2018).

### Linking functional data analysis with deep learning

Regression-based deep learning approaches enable the possibility of mapping image features to continuous numeric data such as plant counting (Ribera et al., 2017), leaf counting (Xie et al., 2023), and plant age (Ubbens and Stavness, 2017). However, in the case of most regression tasks, models map features to output values at fixed points in time. Linking functional traits such as senescence progression requires inputting data into a CNN in a manner that embeds the temporal dimension, which was achieved in this study by concatenating 3-channel RGB images of single plants along the third dimension (time) to create tensors with more than 3 channels (“image sandwiches”). With this method, the data set from the standpoint of input images was reduced from the number of UAS flights × the number of single plants down to the number of single plants while holding the amount of spectral data constant.

FPCA of time-series senescence progression effectively reduced the dimensionality of temporal senescence data, with the first principal component (FPC1) explaining between 83.5% (RCC, Fig. **4b**) to 90.2% (VSR, Fig. **4a**) of temporal variation for combined data from E1 and E2. Variance decomposition of the FPC1 scores by analysis of variance (ANOVA) for VSR and index values for RCC and TNDGR revealed that between 29.9% to 36.8% of temporal variation was attributable to genotype (Fig. **6**). Importantly, this indicates that the FPC1 values carry genotype-level variation through the dimension reduction process. In addition, ANOVA of E1 FPC1 values output by CNN regression indicated that genetic variance partitioning was higher while repeatability values improved (VSR, Fig. **6a**) or remained constant (RCC, Fig. **6b**). FPCA revealed distinct modes of senescence trajectories among individual plants (Fig. **5a**) and indicated that differences in senescence trajectories potentially exist among the chromosome substitution lines (CSLs) and chromosome segment substitution lines (CSSLs) (Fig. **5b**). FPCA also reduces the curse of dimensionality, the “large *p* small *n* problem”, where the number of predictors (*p*) exceeds the number of samples in a study (*n*), posing problems for conventional statistical analysis.

### CNN regression model performance

Learning rate exerted outsized effects on performance of models M1, M2, M4, and M5 but scarcely affected the TNDGR models (M3 and M6, Fig. **7a**). Models M1-M3 and M4-M6 explored the same hyperparameters during optimization (Table **2**) except in the case of M1-M3, where the activation function for the first of two dense layers was always set to ReLU. This had a minimal effect on regression of FPC1 scores for VSRs and TNDGR but affected RCC, where the R^2^ of M2 was ∼42.5% higher than M5 (0.619 vs. 0.434). An inherent limitation of this approach is that initializing the hyperparameter search with different starting conditions and a limited number of trials reduces the likelihood that two Optuna studies converge on the same hyperparameters. However, the high computational cost of running each of the six models in this study limited the trials to 250.

The robust R^2^ values for unseen validation TSIs, especially for VSR FPC1 values (0.857 for M1, 0.886 for M4) demonstrated that both models effectively mapped senescence-related features in the temporally concatenated TSIs. This revealed CNNs can map image features to regression targets with embedded temporal information, thereby uncovering aspects of the organism’s life history as opposed to information about one time point. Saliency maps for M1/M4 confirmed this (Fig. **S1,2**). Despite strong correlations (RCC, *R* = 0.75, TNDGR, *R* = -0.81) with visual senescence ratings (Fig. **3b,c**) and strong (R^2^ = 0.619, 743, and 0.745 for M2, M3, and M6) or moderate R^2^ values (R^2^ = 0.435 for M5), some maps showed inaccurate activation areas outside plant pixels, indicating sensitivity to weed pressure or lighting differences. However, M2/M5 (RCC) and M3/M6 (TNDGR) saliency maps mostly activated correctly at the plant center (Fig. **S3-6**). The lower performance of these models vs. M1/M4 may be due to the larger scale of FPC1 values for visual ratings. Subtle numeric differences for RCC and TNDGR mapped to significant senescence trajectory differences. This was evident in the clustering patterns observed in FPCA of RCC and TNDGR (Fig. **4b,c**) as compared to visual scores (Fig. **4a**). Moderate clustering in TNDGR (Fig. 4c) may explain its higher performance in M3/6 due to greater separation in FPC1 values for rapid senescence vs. stay-green phenotypes compared to RCC. Overall, the VI-based models demonstrate their potential as empirical replacements for VSRs.

### Future directions

Most studies of leaf senescence have investigated the late stages of the aging process, ignoring the earlier developmental stages and their impacts. Future studies are needed that examine senescence in the context of the organism’s complete life history (Woo et al., 2019), particularly deep learning approaches that can handle dense time-series data. To do so, a migration of methods pertaining to functional data analysis into plant sciences is needed, with excellent reviews such as Morris (2015) and Wang et al. (2016) serving as thorough introductions into these topics. The present article demonstrates a novel merger between FDA and deep learning and highlights that CNN regression models can interpret complex temporal phenomena from unseen images, providing evidence that CNNs will likely continue to demonstrate proficiency in regression of other complex, time-dependent agricultural traits. This aim may be bolstered by implementing pre-trained models, which have demonstrated success in tasks such as classifying tomato diseases (Aquil and Ishak, 2021), rice diseases (Shrivastava et al., 2021), and identifying plant pests (Knyshov et al., 2021), though largely in controlled environments which only conduct SPA. Field studies provide additional scale (more plants/plots can be measured) and relevance (real-world signal-to-noise can be evaluated).

The “image sandwich” analysis pipeline developed in this study (Fig. **1**) is adaptable to quantitative temporal analysis of single-plant images of other phenotypes involving spectral changes and could likely be applied to disease scoring or plant growth rates as estimated by vegetation indices. A considerable amount of time within many research programs is spent wrangling and curating data before analysis is conducted, which delays decision-making. CNNs have the potential to minimize or remove the need to annotate data with features such as spatial, environmental, and temporal data in the case of this study. This contributes to an understanding of the phenome of each plant, comprising the summation of all effects within and on a plant regarding genotype and environment (Murray et al., 2022). CNNs can learn these features from raw images and have the potential to supplement and improve visual selection methods that currently persist in plant sciences.

## Declarations

## Acknowledgements

The authors would like to acknowledge Dr. Robert Vaughn and the undergraduate students in the Stelly research group for their time and dedication in maintaining the field experiment as well as Dr. William Beksi and Md Ahmed Al Muzaddid at the University of Texas, Arlington, for their assistance with preparing images for deep learning regression and insightful discussions. A substantial portion of this research was conducted with the advanced computing resources provided by Texas A&M High Performance Research Computing.

## Competing Interests

The authors declare that they have no competing interests.

## Author Contributions

Data curation and analysis were performed by AJD with support from AA. MAA processed and extracted single-plant images and vegetation indices. NS advised model training and saliency visualization. OGR, HB, SMD, and DMS conceptualized and implemented the field experiment. AJD wrote the first draft of the manuscript, and AA MAA, NS, SMD, SCM, OGR, HB, and DMS each contributed to revising and editing of previous manuscript versions. All authors read and approved the final manuscript.

## Data availability

All of the code used for FPCA, ANOVA, and CNN regression is available at the GitHub repository (Notes **S1**) associated with this manuscript (DeSalvio, 2024): https://github.com/ajdesalvio/cotton-sandwiches. All files necessary to run the scripts, including the raw images, are available in the repository.

## Funding

Financial support for this research has been provided by: USDA–NIFA–AFRI Award Nos. 2020- 68013-32371, and 2021-67013-33915, USDA–NIFA Hatch funds, Texas A&M AgriLife Research, and the Eugene Butler Endowed Chair. AJD was supported by the National Science Foundation (NSF) Graduate Research Fellowship (GRFP). NS was supported by the Molecular and Environmental Plant Sciences (MEPS) graduate program. OGR and SMD were partially supported by Cotton Incorporated Awards 18-201 and 20-724, and NSF Award 1739092.

## NSF Statement

Any opinion, findings, and conclusions or recommendations expressed in this material are those of the authors(s) and do not necessarily reflect the views of the National Science Foundation.

## Authors and Affiliations

Interdisciplinary Graduate Program in Genetics and Genomics (Department of Biochemistry and Biophysics), Texas A&M University, College Station, TX 77843-2128, USA Aaron J. DeSalvio (ORCID: 0000-0003-1818-4699), Serina M. DeSalvio (ORCID: 0000-0003- 3741-7144)

Department of Soil and Crop Sciences, Texas A&M University, Agronomy Field Lab 110/111, College Station, TX, 77843, USA

Alper Adak (ORCID: 0000-0002-2737-8041), Mustafa A. Arik, Seth C. Murray (ORCID: 0000- 0002-2960-8226), Oriana García-Ramos (ORCID: 0000-0001-5399-7949), Himabindhu Badavath, David M. Stelly (ORCID: 0000-0002-3468-4119)

Interdisciplinary Graduate Program in Molecular and Environmental Plant Sciences (Department of Soil and Crop Sciences), Texas A&M University, College Station, TX 77843-2474, USA Nicholas R. Shepard (ORCID: 0009-0001-3688-6259)

## List of abbreviations

ANOVA: analysis of variance CNN: convolutional neural network CSL: chromosome substitution line
CSSL: chromosome segment substitution line DAT: days after transplanting
FDA: functional data analysis
FHTP: field-based high-throughput phenotyping FPCA: functional principal component analysis MAE: mean absolute error
MAPE: mean absolute percentage error MSE: mean squared error
ReLU: rectified linear unit RMSE: root mean squared error SPA: single-plant analysis
TSI: time-series image, “image sandwich” UAS: unoccupied aerial system, drone VI: vegetation index
VSR(s): visual senescence rating(s)

## Supporting Information (brief legends) Notes S1. GitHub repository

**Methods S1**. High-throughput phenotyping of cotton experiments.

**Methods S2**. Single-plant shapefile generation. **Methods S3**. Single-plant temporal image extraction. **Methods S4**. Optuna hyperparameter optimization.

**Methods S5**. Model explainability / saliency map generation.

**Table S1.** Vegetation indices used in this study are presented along with respective references.

**Table S2.** Optimal hyperparameters after the conclusion of 250 Optuna trials are presented for M1 and M4.

**Table S3.** Optimal hyperparameters after the conclusion of 250 Optuna trials are presented for M2 and M5.

**Table S4.** Optimal hyperparameters after the conclusion of 250 Optuna trials are presented for M3 and M6.

**Table S5.** Model summary after using the Keras summary() function is presented for M1 (visual scores).

**Table S6.** Model summary after using the Keras summary() function is presented for M4 (visual scores).

**Table S7.** Model summary after using the Keras summary() function is presented for M2 (RCC).

**Table S8.** Model summary after using the Keras summary() function is presented for M5 (RCC).

**Table S9.** Model summary after using the Keras summary() function is presented for M3 (TNDGR).

**Table S10.** Model summary after using the Keras summary() function is presented for M6 (TNDGR).

**Fig. S1.** Activation maps are presented for M1 (visual senescence scores).

**Fig. S2**. Activation maps are presented for M4 (visual senescence scores).

**Fig. S3**. Activation maps are presented for M2 (RCC).

**Fig. S4.** Activation maps are presented for M5 (RCC).

**Fig. S5**. Activation maps are presented for M3 (TNDGR).

**Fig. S6**. Activation maps are presented for M6 (TNDGR).

## Notes

https://github.com/ajdesalvio/cotton-sandwiches

### Competing Interest Statement

The authors have declared no competing interest.

