## Supplementary figures and images for "Temporal Image Sandwiches Enable Link between Functional Data Analysis and Deep Learning for Single-Plant Cotton Senescence"

### Figure S1

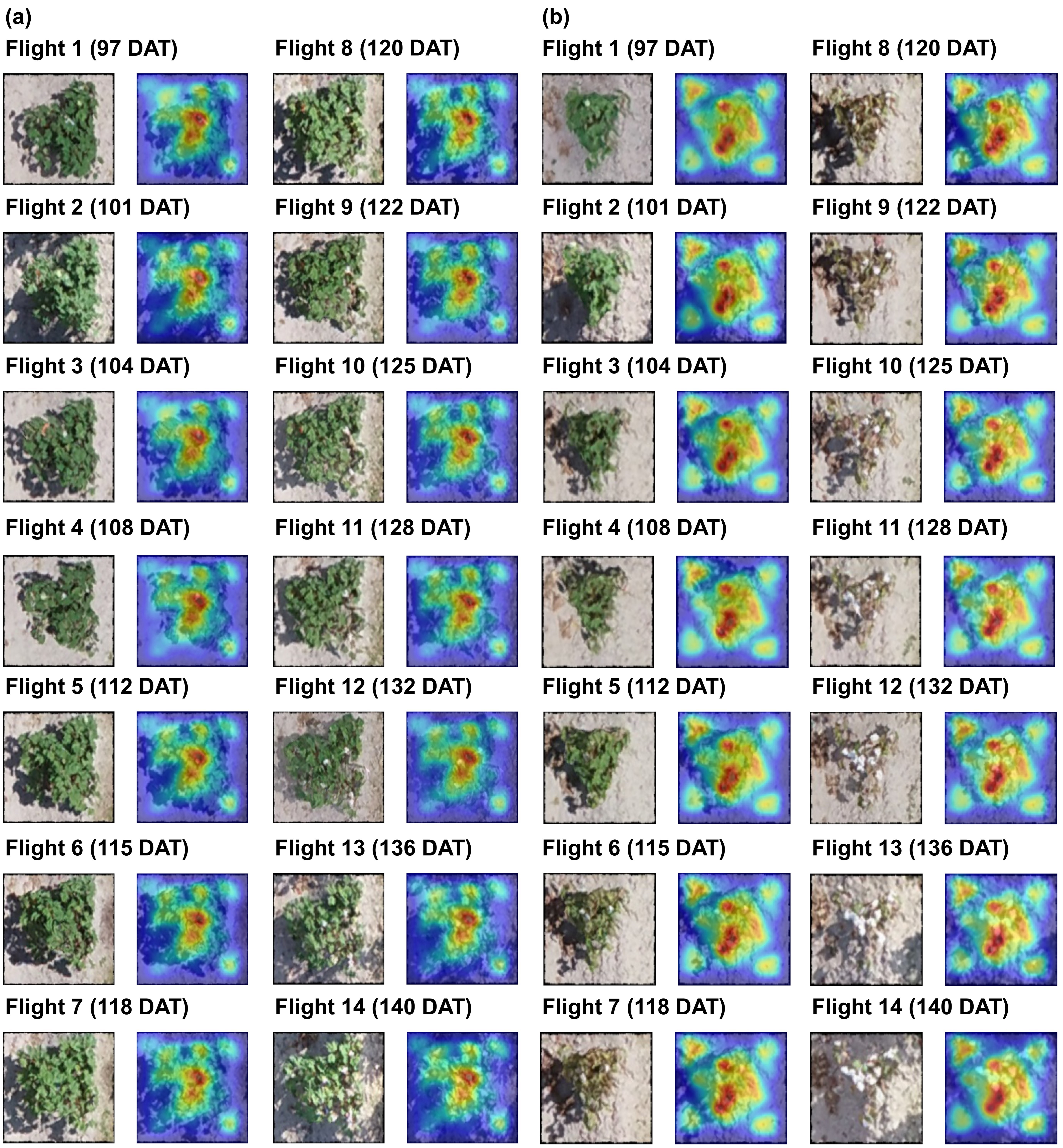

### Figure S2

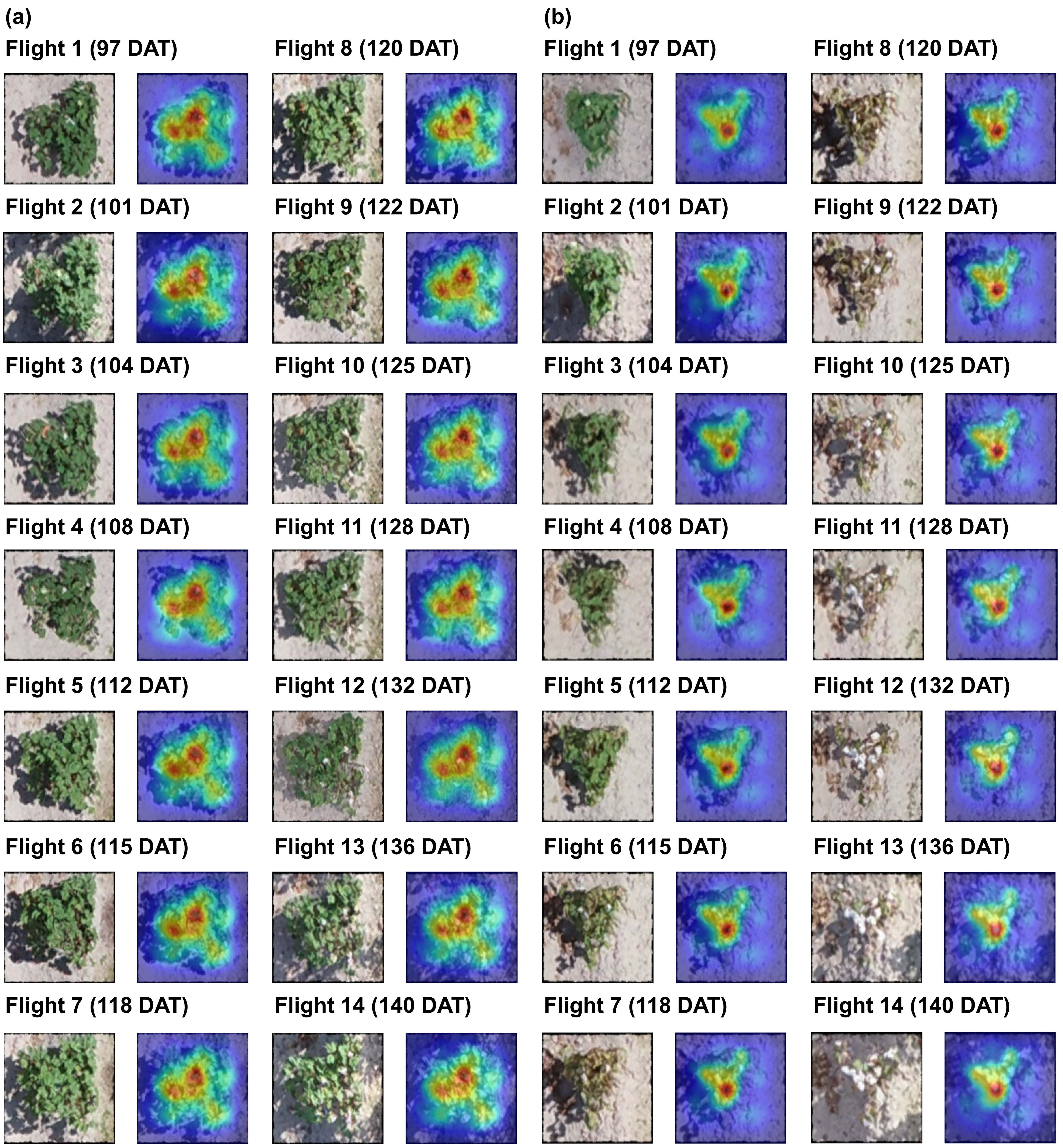

### Figure S3

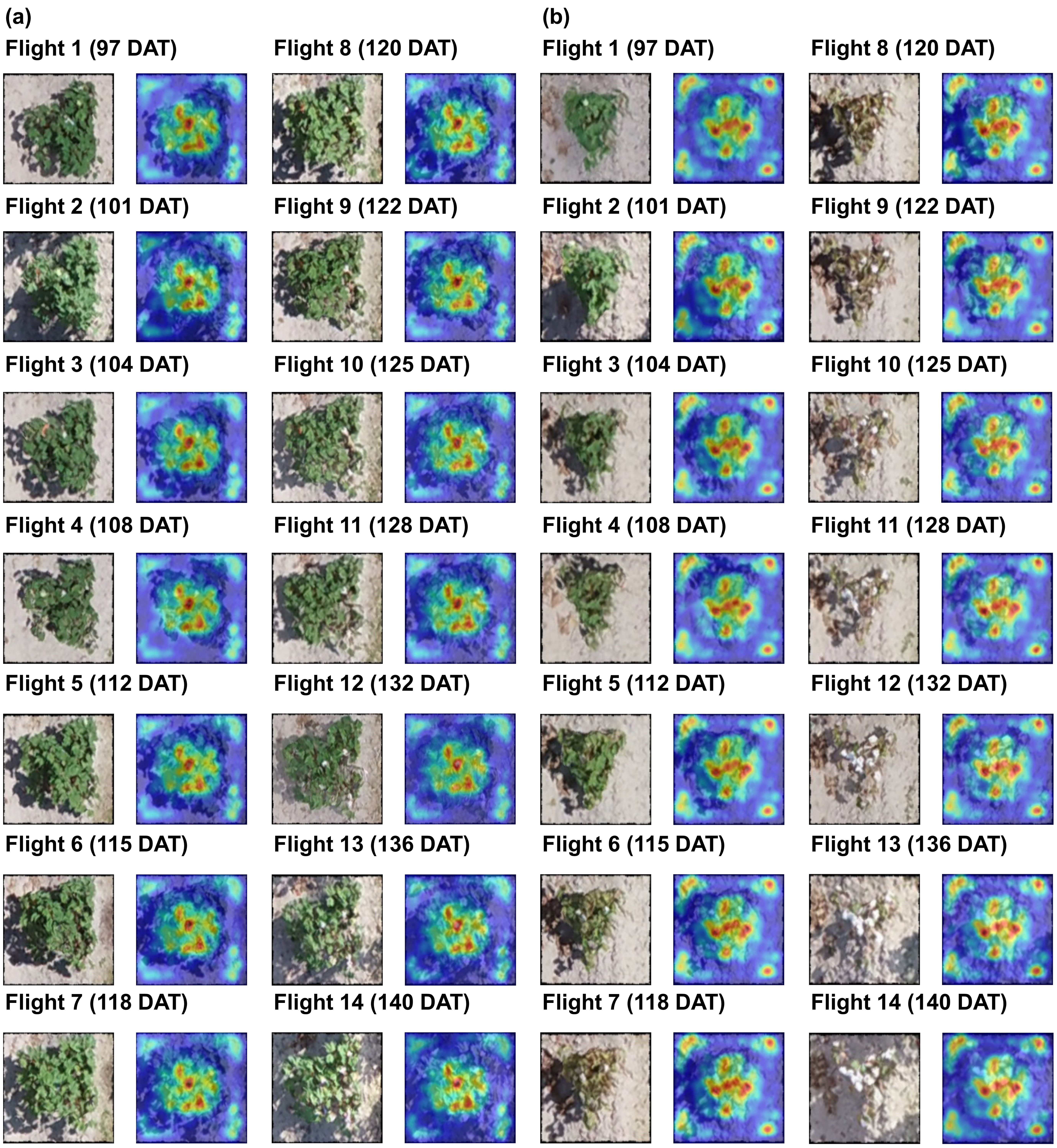

### Figure S4

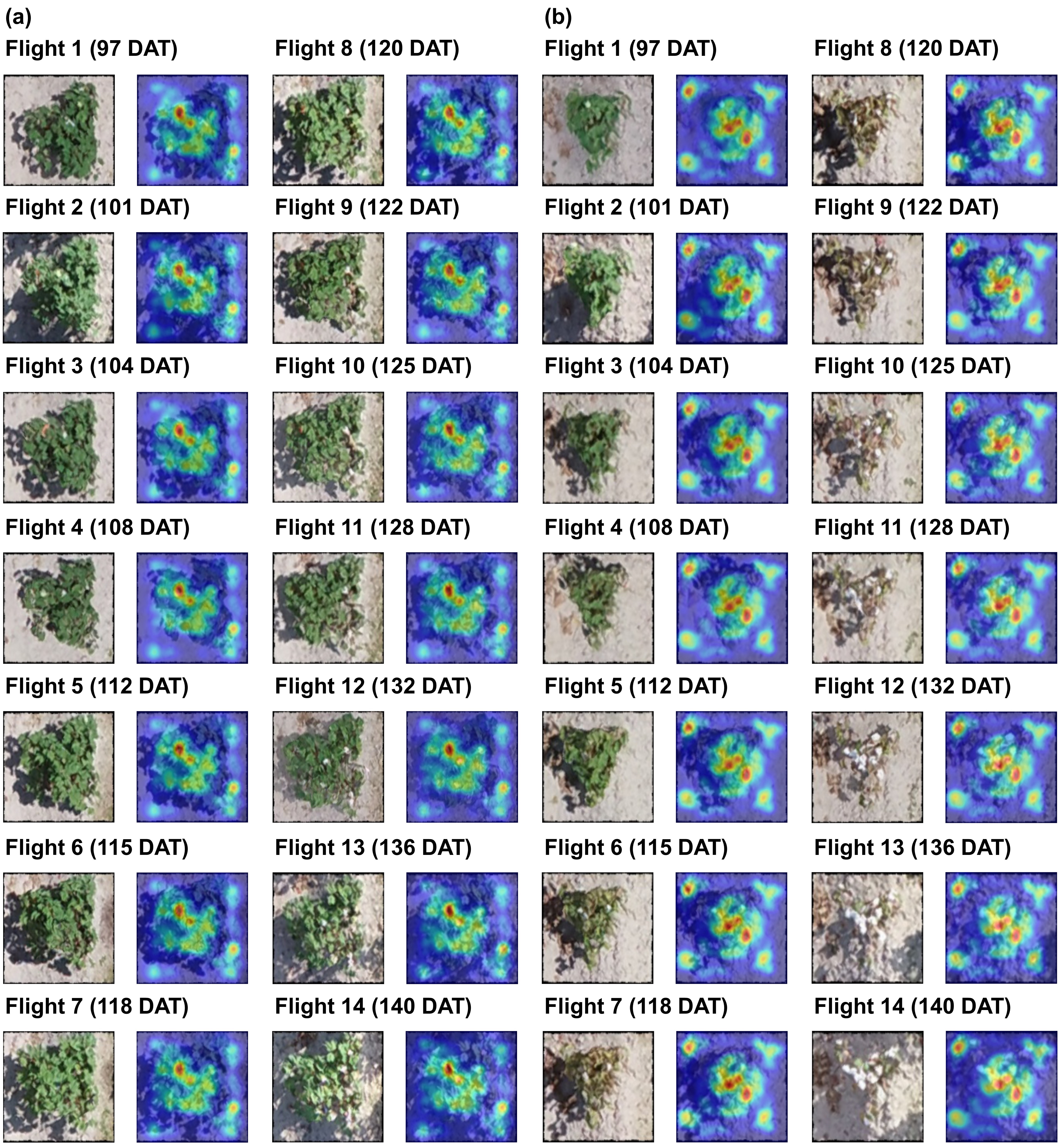

### Figure S5

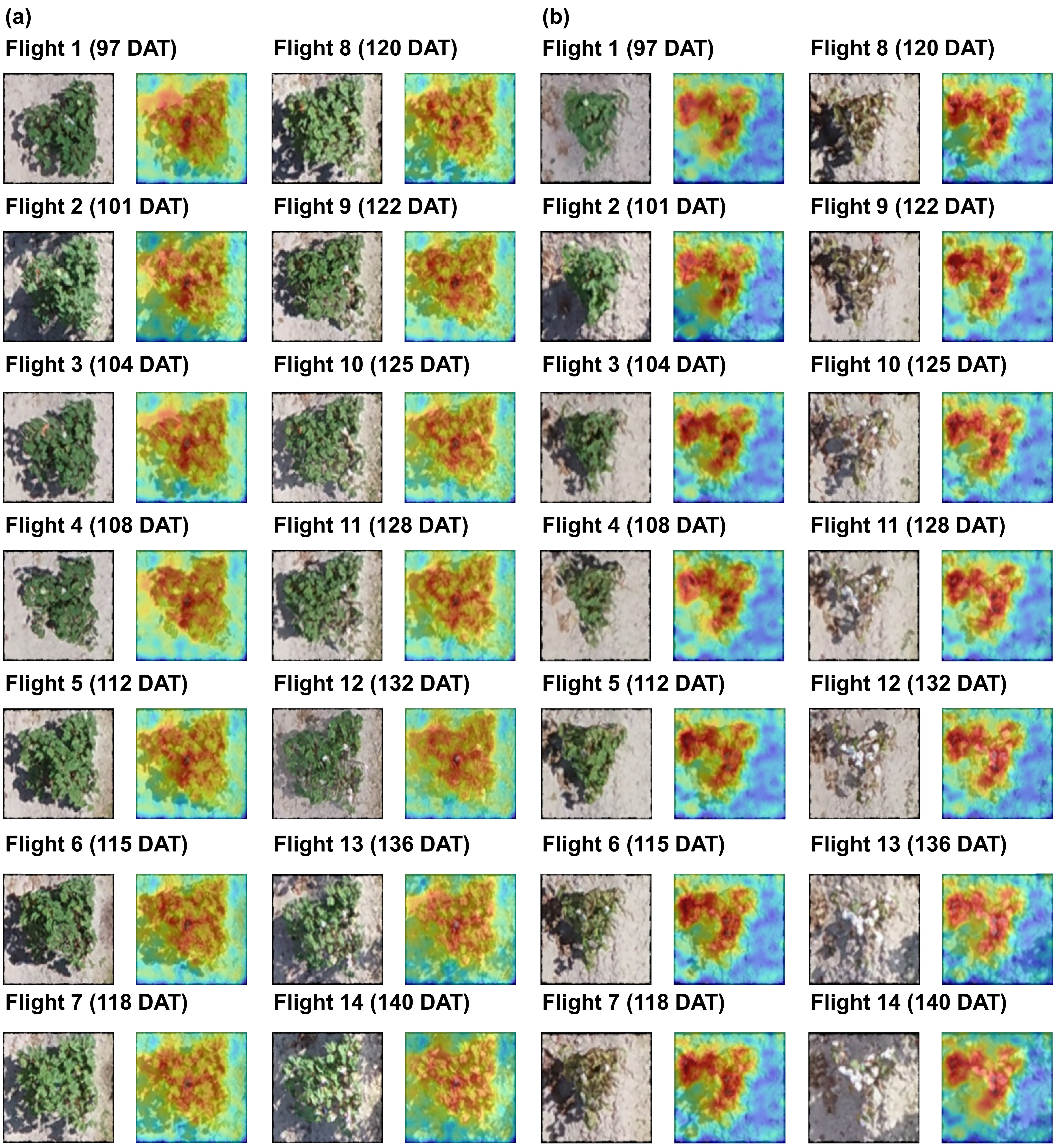

### Figure S6

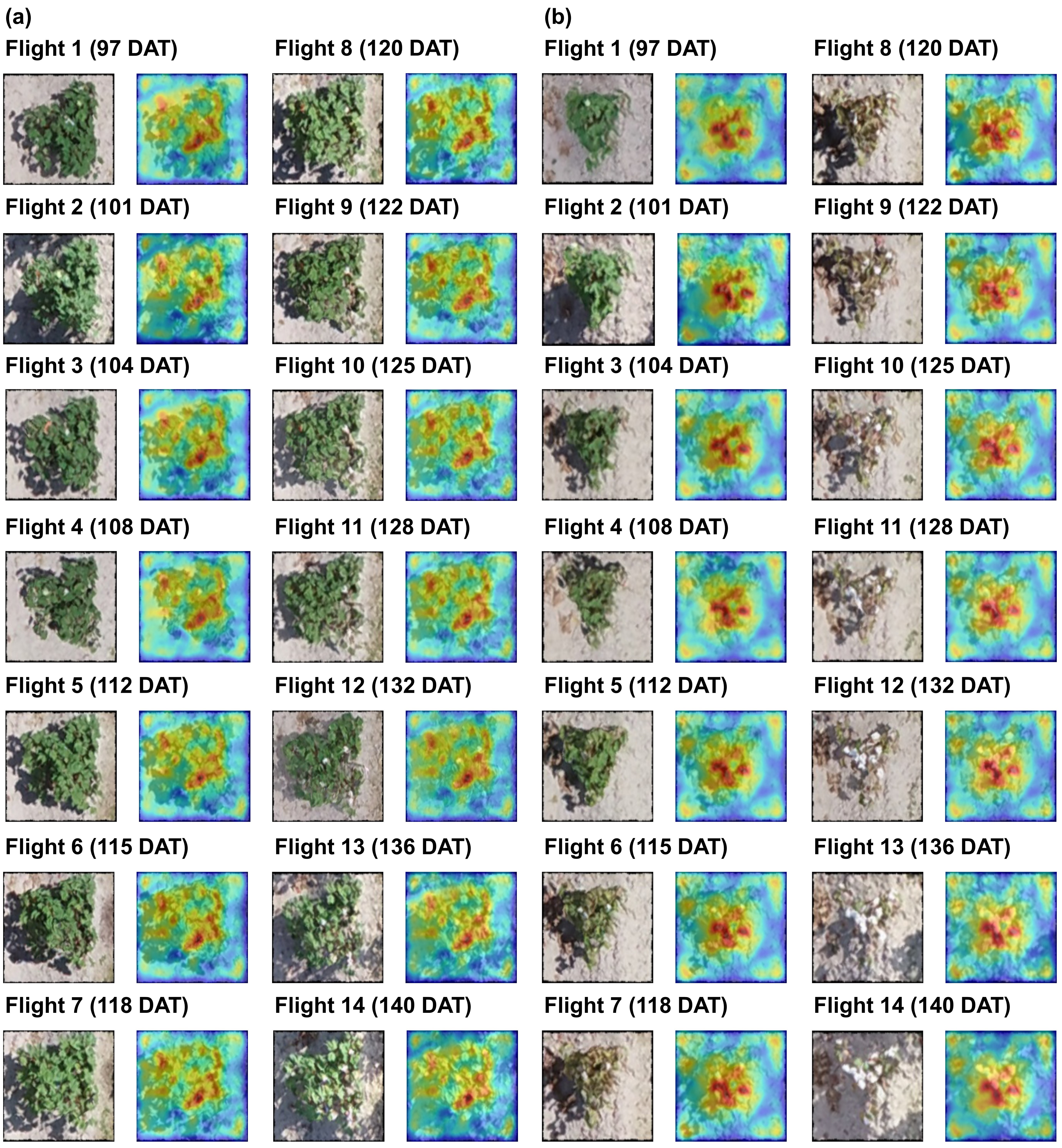
